# Telomeric chromosome ends are highly mobile and behave like free double-strand DNA breaks

**DOI:** 10.1101/720821

**Authors:** Mathias Toulouze, Assaf Amitai, Ofir Shukron, David Holcman, Karine Dubrana

## Abstract

Chromosome organization and dynamics are critical for DNA transactions, including gene expression, replication, and DNA repair. In yeast, the chromosomes are anchored through their centromeres to the spindle pole body, and their telomeres are grouped into clusters at the nuclear periphery, constraining chromosome mobility. Here, we have used experimental and computational approaches to study the effects of chromosome-nuclear envelope (NE) attachments on the dynamics of *S. cerevisiae* chromosomes. We found that although centromere proximal loci were, as predicted, more dynamically constrained than distal loci, telomeres were highly mobile, even when positioned at the nuclear periphery. Polymer modeling indicated that polymer ends are intrinsically more mobile than internal sites. We tested this model by measuring the mobility of a double strand break (DSB) end within a chromosome arm. Upon separation of the DSB ends, their mobility significantly increased. Altogether, our results reveal that telomeres behave as highly mobile polymer ends, despite interactions with the nuclear membrane.

## Introduction

Genome organization is key in controlling genomic functions such as gene expression, DNA damage repair and replication^1–4^. The arrangement of the genome in the nucleus has been described from analysis of loci in population of cells, revealing preferential positions relative to fixed nuclear structures, such as the nuclear envelope (NE) and nucleolus. Chromatin contacts within and between chromosomes and with nuclear structures determine how the genome is folded, which has consequences for its function^5–9^. However how these positions change dynamically in single cells is not fully understood.

The 16 chromosomes in the budding yeast *S. cerevisiae* are bound to the NE through the several interactions. Their point centromeres bind the kinetochores attached to the spindle pole body (SPB), which is embedded into the NE along the whole cell cycle^10–12^. During interphase yeast chromosomes are also tethered to the NE via their telomeres through several pathways involving YKU, Sir4, Esc1, Csm4 Mps3 and the nuclear pore complex (NPC)^13–21^. In addition, Telomeres cluster into 4-6 perinuclear foci at the NE, which favours transcriptional repression mediated by the SIR complex and limits SIR access to other sites of the genome^22–24^. Telomere tethering in S phase contributes also to telomerase control and suppresses recombination among telomere repeats^20,25^. Several intrachromosomal loci, mostly corresponding to either highly transcribed or inducible genes when activated, interact with the NPC imbedded in the NE^26–32^, an interaction that may provide additional layer of regulation and promote optimal gene expression^27,28,33^.

In all organisms genome arrangement varies from cell to cell, and over time, within single cells, due to chromatin motion (visualized using fluorescent repressor operator system [FROS] tagging of single loci and followed at milliseconds time resolution). Chromatin exhibits anomalous subdiffusion that fluctuates with ATP and temperature^26,34–38^. Mobility has been shown to be constrained by the NE and locus position along the chromosome, for example proximity to centromeres or telomeres^2,16,39,40^. Experimental and modelling indicate that the apparent dynamic properties of the chromatin polymer are dictated by these tether points. Centromere tethering strongly restrains motion^2,41,42^ but the consequences of telomere anchoring are less clear. Telomere are more mobile than centromeres^2,41,42^, showing frequent oscillations between the nuclear centre and the nuclear periphery with higher mobility in the nucleoplasm, but with radial motion strongly constrained^16,40,43^. In addition to the constraint arising from tethering, difference in mobility can arise from the crowding of chromosomes chains around the centromeres^44^. A further source of drag on chromatin diffusion comes from the contiguity of the chromatin fibre itself. An extrachromosomal ring excised from a chromosome moves twice faster as the liner parent chromosome and can explore the entire nucleus^45^. While chromatin mobility in the interphase nucleus results from its polymeric nature, the nature and crowding of its environment, and protein-protein interaction that can anchor loci to nuclear structures, it remains unclear how these factors combine to impact chromatin dynamics.

Here we have used experimental and computational approaches to study the effects of chromosome-NE attachments on the dynamics of *S. cerevisiae* chromosomes. Through the analysis of four parameters describing dynamics^46^, we confirm that tethering to the nuclear periphery of both centromere and telomeres constrain their mobility. We reveal that telomeres are more mobile than previously described, with motions similar to intrachromosomal loci even when their mobility is constrained by tethering to the NE. Using modelling and numerical simulations, we predict that the end of a polymer is intrinsically more mobile than an internal site of the polymer. Testing this assumption experimentally, by measuring the mobility of a chromosome extremity following the induction of a double-strand break, we show that free DSB ends have increased mobility. Altogether, our results reveal that the telomeres behave as highly mobile polymer extremities, despite interacting with the NE through protein-protein interactions.

## Results

We examined the dynamics of genomic loci in budding yeast using FROS tags positioned at different position along three chromosomes (Chr III, VI and VII). Lac operators were inserted at 12, 33, 50, 86 and 116 kb from the centromere, visualized by GFP-LacI fusion binding, tracked in live cells over 2 min at 330 ms intervals, and loci positions were measured at subpixels resolution in three dimensions (Figure 1 and S1). Since *LacO* arrays tightly bound by LacI act as proto-silencers, we used a low affinity LacI** protein to minimise potential effects on chromatin structure^47^. DNA dynamics has been shown to vary significantly across the cell cycle^40^. Thus, to avoid data averaging effects due to cell-cycle dependent motion, we selected non-budded G1 phase cells for locus tracking. We analysed the mobility of the loci relative to the telomeres and centromeres by systematically comparing four dynamic biophysical parameters: (i) the length of confinement of the trajectory (Lc) defined by the standard deviation of the locus position, with respect to its mean averaged over time; (ii) the tethering constant (Kc) that estimates the local interaction acting on the locus; (iii) the diffusion coefficient (D) that accounts for local crowding; and (iv) the anomalous exponent, alpha, indicating the nature of the diffusion (see^46^ for calculation details).

**Figure 1:**
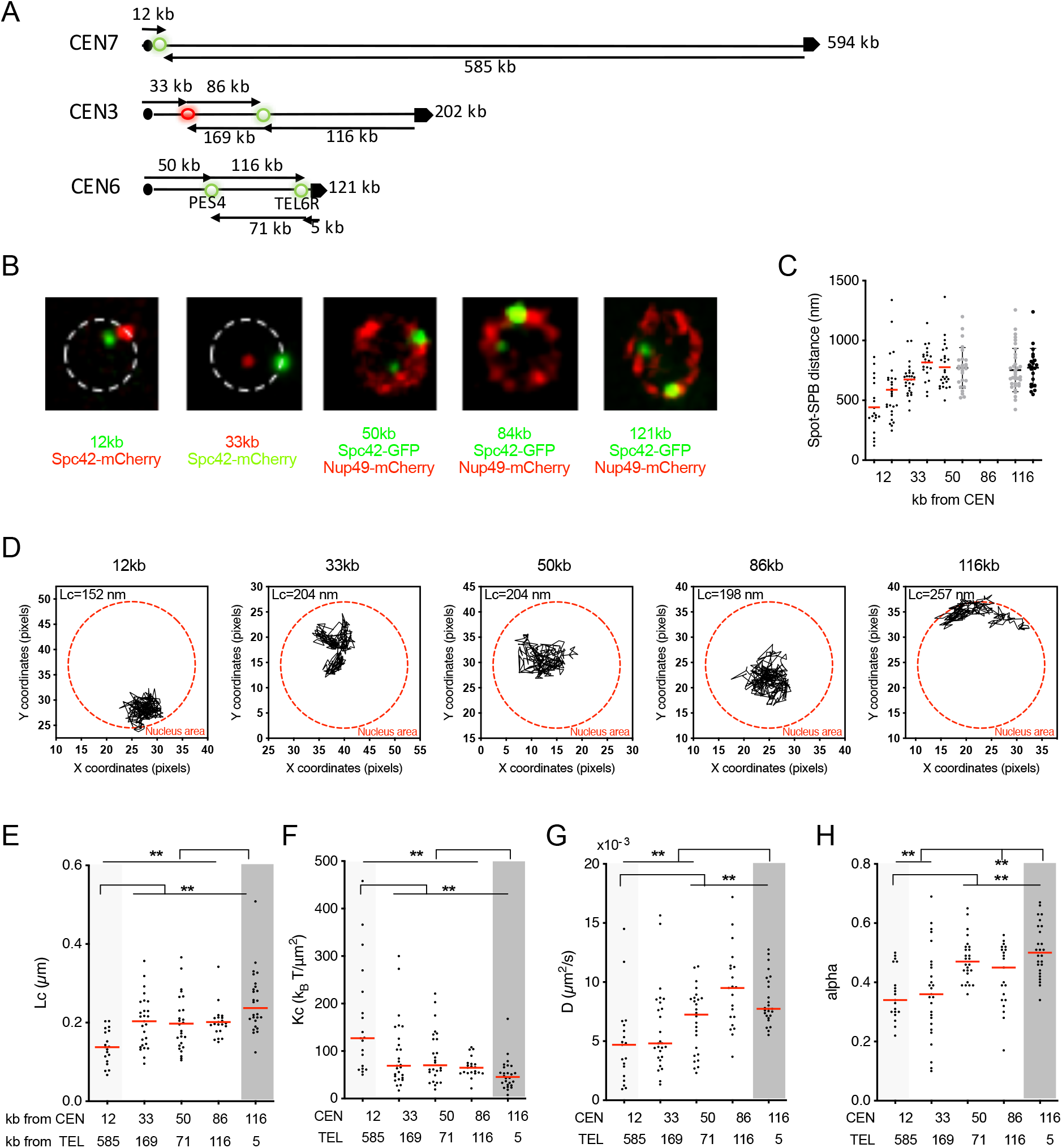
Differential centromere and telomere mobility. A) Chromosomal map of the tagged loci on the right arms of chromosome VII, III and VI. Black dots represent centromeres and black arrows telomeres. The tagged loci are represented with green or red dots depending on their labelling with LacI-GFP or lacI-mCherry respectively. B) Examples of images of LacO/LacI-GFP in the strains used in this study. The tagged loci and their distance from the centromere are indicated. LacO/LacI-GFP is the weakest green signal; the strongest green signal is Spc42-GFP; Nup49-mCherry is in red. C) Plot of the LacO/LacI-GFP spot distance to the SPB in nm for each movie analysed. Red bar, distribution median. Black stars indicate statistical differences (* = p<0,05; ** = p<0,01; *** = p<0,005; **** = p<0,001). D) Representative trajectories of the indicated loci during ∼ 120 s (∆t=300 ms). Lc of the corresponding movie is indicated. D-F) Distribution of the indicated parameters for the indicated loci. Red bar, distribution median. Black stars indicate statistical differences (* = p<0,05; ** = p<0,01; *** = p<0,005; **** = p<0,001). Centromeric and telomeric loci are shaded in light grey and dark grey respectively.

The nucleus itself exhibits both spatial translation and rotation that translate into parasitical movement of the tracked loci^46^. To estimate the effect of nuclear movements followed by SPB motion, we plotted the four parameters Lc, Kc, D and alpha with respect to the mean confinement length of the SPB (Lc^SPB^) for each cell. In cells with Lc^SPB^ higher than 200 nm the confinement length of the locus was systematically above the median Lc of the population and the tethering constant below the median of the population showing that nuclear movement has a strong impact on these two parameters (Figure S1). The diffusion coefficient D and the anomalous exponent alpha were not clearly affected. These results are consistent with polymer model simulations where the frequency and amplitude of the oscillation of the tethering point of a polymer confined in the nucleus affect the values of Lc, Kc, D but not alpha of nearby loci (Figure S2, see also Figure S3 for single simulation runs). The influence of the boundary oscillation was no longer detected 10 monomers from the tethering point (in a total of 100 monomers per chromosome), as measured by the four parameters Lc, kc, Dc and alpha (Figure S2). We conclude that the oscillation of the boundary could influence centromere and telomere dynamics, but not the dynamics of loci located at a genomic distance less than 10% of the total chromosome length from the anchor point. Thus, to eliminate parasitical nuclear movement, we considered only those cells with limited SPB movement (Lc^SPB^ < 200 nm; Figure S1, see Material and methods).

### Differential centromere and telomere mobility

The mean distance between the imaged locus and the SPB increased with the distance between the locus and the centromere, ranging from 442 nm for *CEN7*, located 10 kb from the centromere, to 816 nm for *MAT*, located 86 kb the from centromere (Figure 1C), confirming that the centromere constraints intranuclear position. However, the telomere proximal locus *TEL6R*, at 116 kb from the centromere, and the intrachromosomal *MAT* locus, at 86 kb from the centromere, exhibited similar mean distances from the SPB. Thus, additional constraints act on telomeres, likely reflecting the tethering to the NE. Examination of the trajectories after projection onto a single x-y plane during time lapses showed that *TEL6R* traces a crescent-shaped track close to the NE whereas intrachromosomal loci sample a large fraction of the nucleoplasm (Figure 1D). The track of the centromere proximal locus *CEN7* is restrained close to the NE, consistent with its anchoring to the SPB, but unlike *TEL6R* it does not follow the NE. Accordingly, *CEN7* exhibited the lowest spatial constraint with Lc^12kb^= 0.14 ± 0.04 µm. In agreement with previous data, Lc increased rapidly as the distance from the centromere increased reaching a plateau such that it did not differ statistically for loci at 50 kb and 86 kb (Figure 1E;^2^). This increase in motion correlates with a substantial decrease in the strength of the tethering forces as the distance from the centromere increases (Figure 1F). These results indicate that centromere proximal loci are subjected to strong local interactions which impacts decrease along the chromosome. These local interactions are likely caused by tethering of the centromeres to the SPB, disruption of which increases motion^2^. Whereas Lc and Kc did not differ between loci located more than 33kb away from the centromere, the coefficients of diffusion (D) were similar for the 12kb and 33kb centromere proximal loci but significantly increased for loci more than 50kb away from the centromere (Figure 1G). As previously described, all the loci analysed showed sub-diffusive motion, with mean values for alpha ranging from alpha^12kb^=0.36 to alpha^116kb^= 0.51 and a large cell-to-cell variation^19,26,34,35,37,38^. As for D, alpha values were low and indistinguishable for the two centromere proximal loci at 12 and 33 kb and significantly higher for loci located beyond 50 kb (Figure 1H).

These results confirm that centromere attachment to the SPB constrain mobility of proximal loci (within 10 kb)^2^. They also reveal that the coefficient of diffusion and the anomalous exponent are also constrained by centromere proximity, at larger distances (up to 33kb).

Our results did not reveal any of the expected characteristics for NE tethered telomeres. The 116kb telomere proximal locus *TEL6R* was more mobile than more centromere proximal loci and exhibited the highest Lc and the lowest Kc (Figure 1E-H). The associated diffusion coefficient D^116kb^ was only slightly lower than that of the 86 kb intrachromosomal locus and significantly higher than that of the 50 kb locus, located in the middle of the same chromosome (Figure 1F). The telomeric locus also exhibited an anomalous exponent that was significantly higher than that of the 86 kb intrachromosomal locus and of the 12 kb and 33 kb centromere tethered loci (Figure 1G). These high alpha values are unlikely to be caused by nuclear rotations or oscillation as we restricted our analysis to nuclei with low Lc^SPB^ and do not observe them for centromeric proximal loci anchored to the SPB. Rather they seem to reflect the behaviour of subtelomeric loci.

### Telomeres are highly mobile despite perinuclear localization

To further understand the high mobility of telomeres, we analysed individual *TEL6R* movies, which revealed that telomeres dynamically switched from peripheral to more internal positions in G1 cells (Figure 2A). Telomeres spending most of the time close to the nuclear membrane were more constrained than those more centrally located, yet their length of constraint Lc was above 200 nm similar to intrachromosomal loci (Figure 2A). This high mobility could result from the release of the tethering constraint, as shown for centromeres whose motion increases upon detachment^2^ and for telomeres located in a central position or in mutants that disrupt their peripheral anchoring ^16,34,39^. Alternatively, the change in position by itself, which changes the environment could account for the higher mobility of internal telomeres.

**Figure 2:**
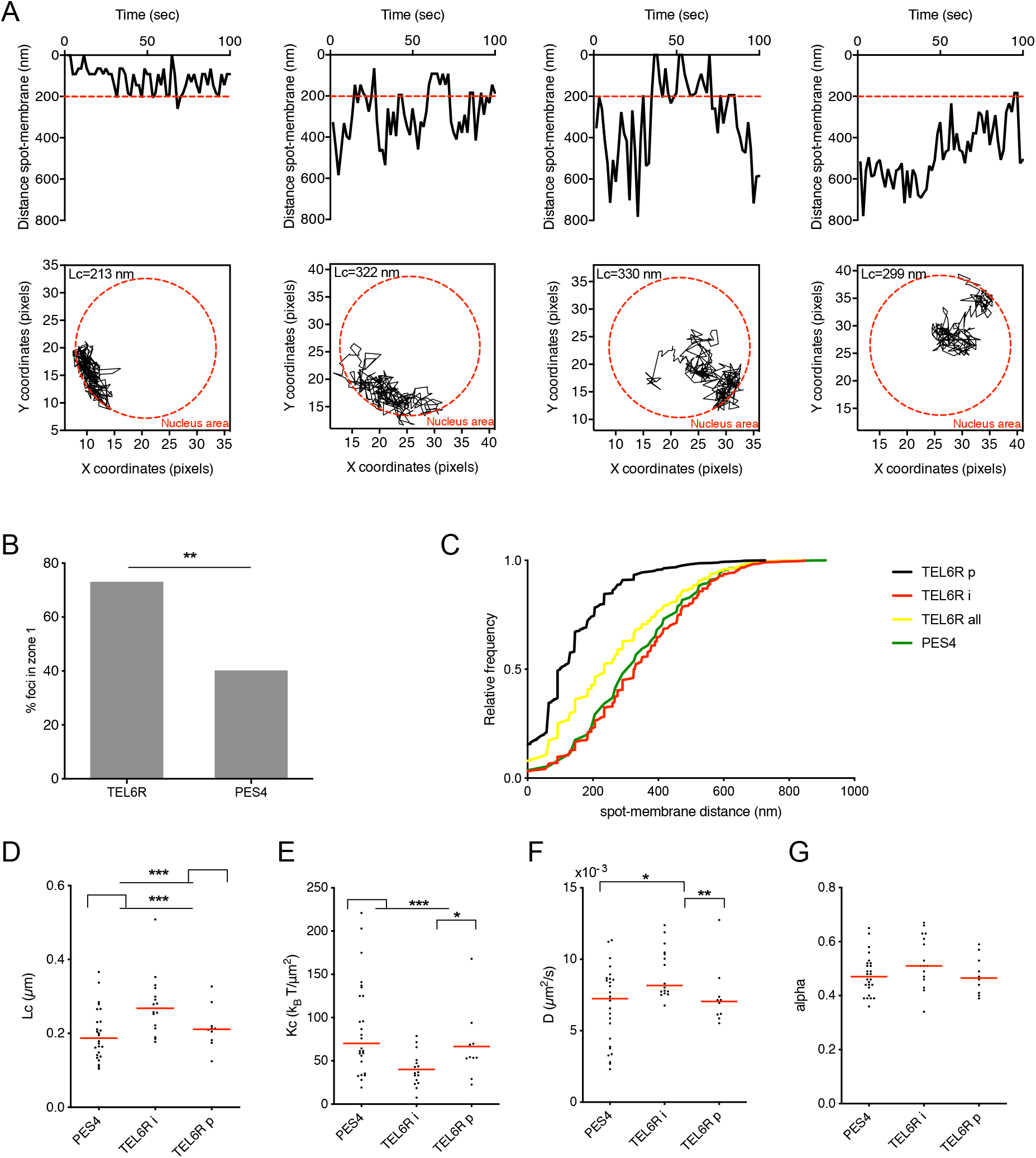
Motion of internal and peripheral telomeres. A) Plot of the spot-to-membrane distance along four *LacO*-tagged *TEL6R* representative movies (top panels) and corresponding trajectories during ∼ 120 s (∆t=300 ms) (bottom panels). B) Position of *LacO*-tagged PES4 and *TEL6R* in *WT* and *∆ku80* strains was determined in snap shot images. Locus position is scored relative to the nuclear periphery stained by Nup49-mCherry in its plane focus on population of cells (see Material and methods). Bars represent the percentage of cells with loci located less than 200 nm from the nuclear periphery. **= significantly non-random distribution on the basis of a 95% confidence interval of a proportional test between the experimental distribution. C) Cumulative distribution function for the distance of the spot to the nuclear envelope (NE) in all the movies analysed for *TEL6R* (*TEL6R* all, yellow line) and *PES4* (*PES4*, green line). The cumulative distribution of *TEL6R* movies with a mean distance to the NE below 200 nm (*TEL6R* peripheral, red) and above 200 nm (*TEL6R* internal, black) are shown. D-G) Distribution of the extracted parameters of the indicated loci. Red bar, distribution mean. Black stars indicate statistical differences (* = p<0,05; ** = p<0,01; *** = p<0,005; **** = p<0,001).

To test whether the higher mobility we observed was due to the number of cells with non-peripheral telomeres in our strain, we estimated *TEL6R* position relative to the nuclear periphery in snapshot images (Figure 2B). The shortest distance between the locus and the NE was measured. Signals located within 200 nm were considered as peripheral. In agreement with previous results^16,40^, *TEL6R* was enriched at the nuclear periphery in 70% of the cells, whereas the intrachromosomal *PES4* locus, located on the same chromosome, exhibited a more random positioning and was located at the nuclear periphery in 40% of the cells (Figure 2B). We quantified the position of *TEL6R* and *PES4* relative to the nuclear periphery in all movies, analysing the cumulative frequency of spot-membrane distances, and found that *TEL6R* is globally closer to the NE than *PES4* (Figure 2C, Figure S3). Contrary to the observations in snap shot images, TEL6R was located in the 200 nm outermost zone of the nucleus in less than 40% of the time frames, indicating that the movies analysed are biased toward internal telomeres. To correct for this bias, we analysed separately movies in which *TEL6R* is mostly peripheral (TEL6Rp), with a mean distance from the periphery below 200 nm, and movies in which *TEL6R* is internally located (TEL6Ri), with a mean distance from the periphery above 200 nm (Figure 2C). We observed that internal telomeres (TEL6Ri) were more mobile than the intrachromosomal locus PES4 on the same chromosome, with higher Lc, Kc and D, revealing that telomere are more mobile than intrachromosomal loci despite moving in the same environment. When located at the nuclear periphery, (TEL6Rp) telomeres are less mobile than internal one (TEL6Ri) and exhibit a higher tethering constraint, consistent with the perinuclear anchoring^14,16^. The diffusion coefficient was also lower for peripheral telomeres with D^TEL6Rp^= 7.38 ± 2.09 10^−3^ µm^2^/s compared to D^TEL6Ri^= 9.05 ± 1.77 10^−3^ µm^2^/s (Figure 2D-E). However, the nature of the diffusion remained unchanged as alpha was not statistically different between TEL6Ri and TEL6Rp (Figure 2G). As the anomalous exponent depends in part on intrinsic polymer properties, this suggests that the chromatin structure of internal and peripheral telomeres is not detectably different. Despite the constraint imposed by perinuclear anchoring, we surprisingly observed that the mobility of the most peripheral telomeres was similar to that observed for the intrachromosomal locus *PES4* on the same chromosome, with none of the dynamic parameters being statistically different between TEL6Rp and *PES4* (Figure 2 D-G).

These results reveal that, on short time scales (2 min movies), telomere motion is constrained by interactions with the NE, yet this constraint is not sufficient to lower telomere mobility below that observed for the intrachromosomal locus PES4.

### Telomeres-perinuclear component interactions constrain mobility

Telomere perinuclear localisation has been shown to rely on redundant pathways involving the inner membrane–binding protein Esc1, Mps3, the NPCs, the telomere-binding SIR and yKu70/80 complexes^16,27^. Since *TEL6R* perinuclear localization has been shown to depend mainly on the yKu70/80 pathway^16,39^, we tracked *TEL6R* motion in cells mutated for *YKU80* (Figure 3A). We confirmed that deletion of *YKU80* abolishes perinuclear localisation of *TEL6R* in snap shot images (Figure 3B). *TEL6R* also adopted an intranuclear localisation in time-lapse series similar to intranuclear telomeres (TEL6Ri; Figure 3C). We analysed *TEL6R* dynamics quantitatively using imaging in WT and *yku80∆* cells comparing Lc, Kc, D and alpha. The four parameters values were not significantly different in *YKU80* deleted cells compared to WT internal telomeres (TELR6Ri; (Figure 3D-G)). However, Lc, Kc, and D of *TEL6R* were significantly different when peripheral WT telomere TEL6Rp was compared to unanchored *TEL6R* in the *ku80∆* mutant (Figure 3D-F). We measured higher Lc and lower tethering force Kc upon loss of anchoring in *yku80∆* cells compared to peripheral telomeres TEL6Rp demonstrating that *TEL6R* can explore a larger space when no longer attached to the NE. It is unlikely that this increased motion results from the modification of the telomere chromatin structure in the *yku80∆* mutant, as analysis of the fraction of *TEL6R* that are located internally also exhibit this trend. We observed no change in the anomalous exponent alpha despite telomere increased motion upon detachment, suggesting that the characteristics of the motion remained unchanged (Figure 3G).

**Figure 3:**
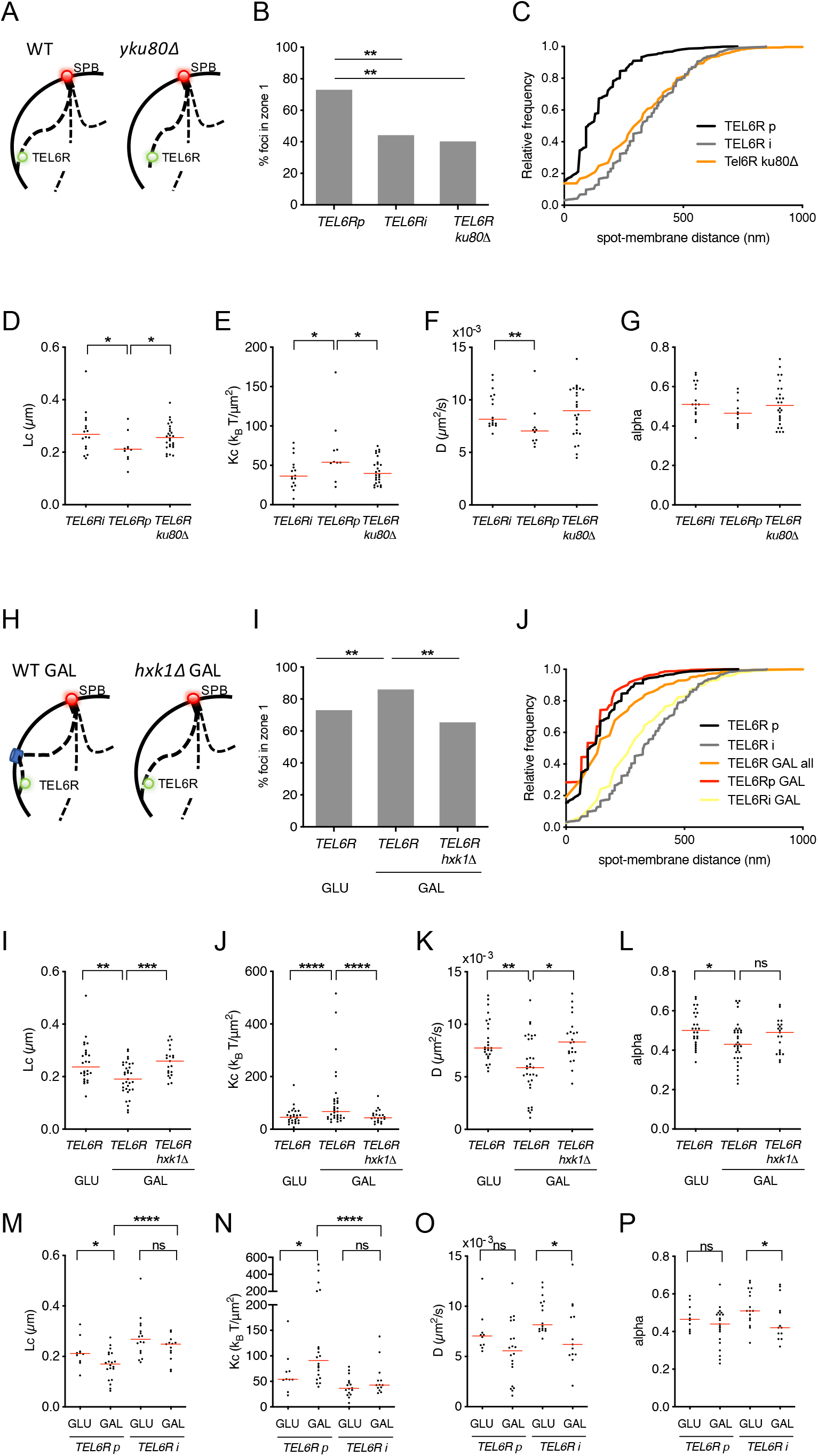
Interactions with the nuclear periphery constraint mobility. A) *TELR6R* loses nuclear membrane anchoring in *ku80∆* mutants. B) Position of *LacO*-tagged internal and peripheral *TEL6R* in *WT* and *ku80∆* strains were determined with respect to the NE as in Figure 2B. C) Cumulative distribution function for the distance of the spot to the NE in all the movies analysed for peripheral and internal *TEL6R* and in *ku80∆* cells. D-G) Distribution of the extracted parameters for the indicated loci and in the strains mentioned. Red bar, distribution mean. Black stars indicate statistical differences (* = p<0,05; ** = p<0,01; *** = p<0,005; **** = p<0,001). H) TEL6R is anchored to both the nuclear pores and the NE in WT grown in galactose. Nuclear pore tethering is lost in hxk1∆ cells grown in galactose but telomere tethering remains. I) Position of *LacO*-tagged *TEL6R* in WT strain grown in glucose (*TEL6R* GLU) and in WT or *hxk1∆* strain grown in galactose (*TEL6R* GAL and *hxk1∆* GAL) was determined with respect to the NE as in figure 2B. J) Cumulative distribution function for the distance of the spot to the NE along all the movies analysed for peripheral and internal *TEL6R* in glucose or in galactose. I-P) Distribution of the extracted parameters of the indicated loci and in the growth conditions mentioned. Red bar, distribution mean. Black stars indicate statistical differences (* = p<0,05; ** = p<0,01; *** = p<0,005; **** = p<0,001).

Altogether, untethering of *TEL6R* upon *YKU80* deletion increased both dynamics and nuclear volume explored indicating that, like centromeres, tethering of telomeres restrains their mobility, although to a lesser extent, further suggesting that the nature and the strength of the tethering influence chromatin dynamics.

### Additional nuclear membrane anchoring points constrain mobility

To get further insights into how tethering affects mobility, we imaged cells in which *TEL6R* was attached to the nuclear membrane through an additional anchor site. Indeed, *TEL6R* relocalises toward the nuclear periphery in galactose medium as a consequence of the transcriptional activation of the nearby *HXK1* gene and its interaction with the NPC^27^. The *HXK1* gene, which is located 1.8 kb from the fluorescent tag on *TEL6R*, is repressed in glucose medium but its transcription is strongly induced in absence of glucose (^48^; Figure 3H). Thus, in WT cells grown in galactose, *TEL6R* will be anchored both to NPC via the *HXK1* gene and to the NE via the telomeres. Growth in galactose induces large scale changes in gene expression and chromosome organisation^32^. As a control, we used a strain deleted for *HXK1* (*hxk1∆*) in which *TEL6R* is only tethered via its telomere. In galactose medium, *TEL6R* shifted toward the nuclear periphery. This shift was lost upon *HXK1* deletion despite growth in galactose medium, consistent with earlier findings (^27^; Figure 3I). The plot of the cumulative frequency of movie spot-membrane distances showed that *TEL6R* is overall close to the NE upon *HXK1* induction, behaving like the most peripheral TEL6Rp in glucose medium (Figure 3J). The dynamics parameters from the doubly tethered *TEL6R* (*TEL6R* GAL; Figure 3 I-L) were significantly different from those observed for *TELR6* with a single tether (*TEL6R* GLU and *TEL6R hxk1∆* GAL; Figure 3I-L). This difference was not due to the change of sugar as TEL6R with a single tether in glucose (TEL6R GLU) and in galactose (*TEL6R hxk1∆* GAL) exhibited equivalent dynamics. Rather, induction of an additional anchor to the NE *per se* increased the tethering constraint, Kc, and concomitantly reduced the length of constraint, Lc (Figure 3I-J). It also decreased the diffusion coefficient, suggesting that the locus encounters a more crowded environment when anchored at the NPC (Figure 3K). The anomalous exponent alpha also decreased upon NPC targeting, indicating that TEL6R explores space in a highly recurrent manner when the locus is flanked by two tethering points (Figure 3L). This result suggests that the anomalous exponent, known to be sensitive to chromatin compaction^46^, can also capture higher order chromatin folding.

We observed two cell populations in galactose, those with peripheral telomeres (TEL6Rp) and those with internal telomeres (TEL6Ri) (Figure 3M-P), similar to TEL6R telomeres in glucose. The diffusion coefficient, D, and the anomalous exponent, alpha, decrease slightly for both peripheral and internal telomeres in galactose medium, although this decrease was only statistically significant for intranuclear telomeres (TEL6Ri) in galactose compared to glucose medium (Figure 3O-P). TEL6Ri Lc and Kc remain unchanged in galactose compared to glucose, in agreement with the internal localisation (Figure 3M-N). This suggests that *HXK1* transcriptional activation, which also occurs in the nuclear interior^27^ significantly modifies the diffusion rate and the nature of the motion independently of tethering effects.

When comparing the motion of telomeres in the most peripheral zone (TELR6Rp) after growth in glucose or in galactose, only Lc and Kc were statistically different. Consistently with analysis of the whole population, peripheral telomeres with an additional anchor are more highly tethered, with increased Kc and decreased Lc values (Figure 3M-N). Kc roughly doubles with the presence of a second anchor point. If we consider that the two tethers are described as two harmonic springs of constant K_1_ and K_2_ acting in parallel, then the resulting spring constant, K0, is given by: 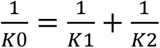. Since the experimental spring constant, Kc, doubles with the addition of one tether, this suggests that K_1_ =K_2_ thus 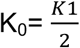 as predicted by the law of parallel springs. This result shows that Kc can quantitatively capture changes in the tethering force as subtle as the addition of a second local tethering point *in vivo*.

### Polymer simulations recapitulates yeast chromatin dynamics

We performed polymer simulations and compared them with our experiments to address the role of centromere and telomere tethering on chromosome dynamics in the interphase nucleus of yeast cells. We used a Rouse polymer model of yeast chromosomes that incorporate various tethering constraints consistent with their organization. We modelled the nucleus boundary as a sphere of radius 1 µm. Chromosomes were tethered via their central monomer (50) to a fixed point on the membrane, simulating CEN-SPB attachment. Monomers 1 and 100 at the extremities, mimicked telomeres. Monomers 2 to 99 were allowed to move inside the 1µm sphere. To test the effect of telomere tethering on dynamics parameters, we computed Lc, Kc, D and alpha in the presence or in the absence of a fixed tether at both monomer 1 and 100 versus when both were allowed to diffuse freely across the surface of the sphere (model parameters are described in supplementary methods subsection 1). The fixed tethers mimicking the centromere-SPB and telomere attachments affected all four parameters in the immediate vicinity of the tethering point, which, based on our model parameters was estimated to be 1% of the polymer length (Figure 4). This is consistent with previous modelling showing that fixed tethering only constrains dynamics of genes less that 10kb from the tethering point^2,42,49^. This is, however only qualitatively consistent with our experimental results, which showed that the coefficient of diffusion, D, and the anomalous exponent, alpha, are affected up to 33 kb from the tethering point (Figure 1G-H). In addition, a fixed tether was not consistent with the high dynamics observed for the *TEL6R* bulk population (Figure 1G-H) nor for the most peripherally located telomeres TEL6Rp (Figure 2F-G). Conversely, simulations with either telomere diffusing on the nuclear membrane, or with telomeres moving freely in the nucleoplasm, agreed well with our experimental data for *TEL6R* (Figure 4).

**Figure 4:**
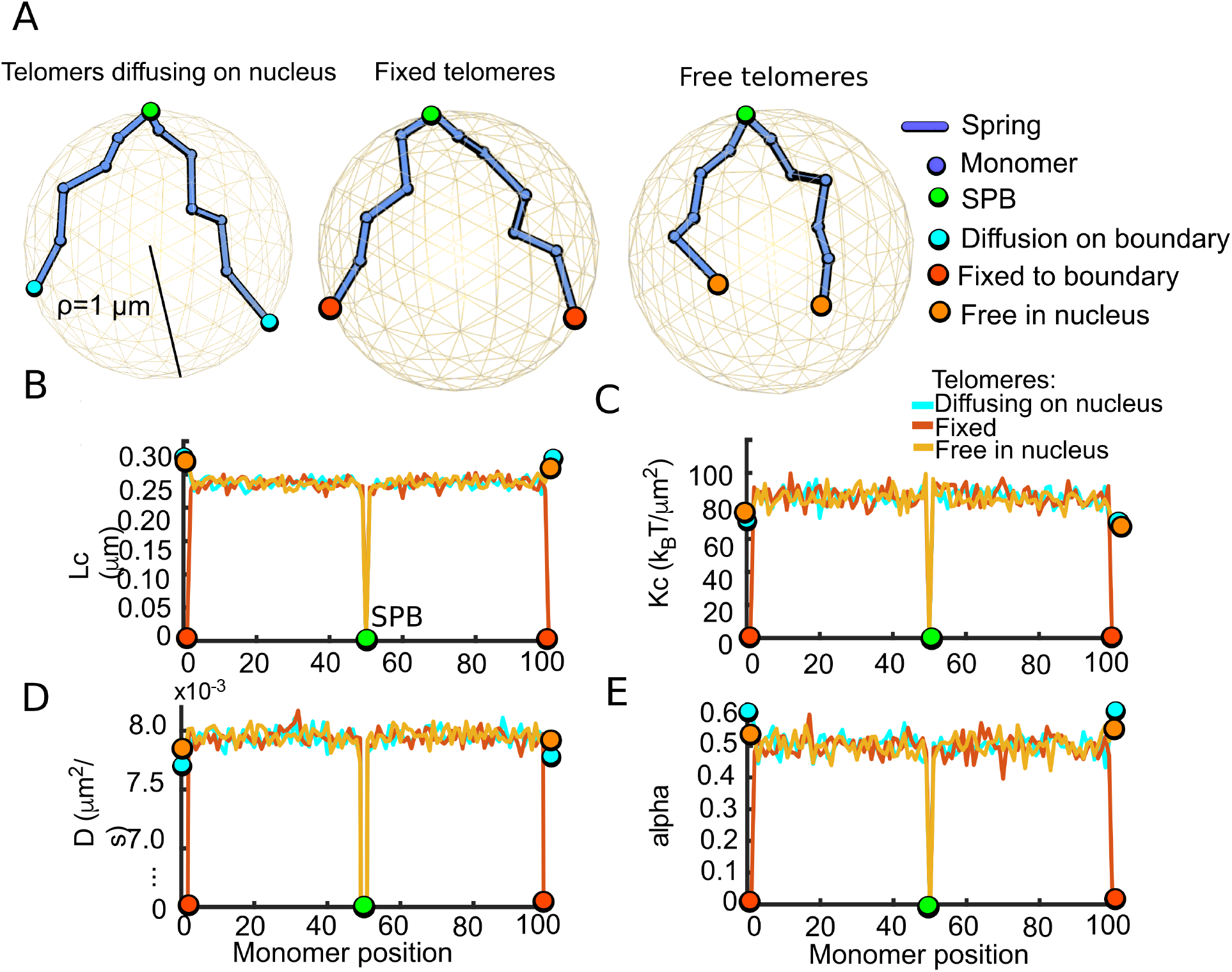
A Rouse polymer model cannot recapitulate chromatin mobility parameters. A) Sketch of Rouse polymer model located inside a reflecting spherical domain of radius= 1µm, containing N = 100 monomers (purple spheres). The end of a polymer (= telomere) is either fixed on the surface or can move by Brownian motion B-E) Four parameters description, simulations of the polymer described in panel A: B) Length of confinement Lc, C) Effective spring constant Kc, D) Diffusion coefficient D, E) Anomalous exponent.

Given the Rouse model did not simulate the chromosome dynamics revealed by the four parameters (Figure 4), we used a modified randomly cross-linked polymer^50^. To account for the inter-chromosomal interactions between pericentromeric regions we added random connectors between monomers centred around monomer 50 (representing the anchored SPB). Connectors were added according to a normal distribution, using an optimal fit to the experimental single particle trajectory data, leading to 60 added connectors (Nc=60) with a variance = 9 (Figure 5A). Using these parameters, we ran the simulations for 60s, with a sampling time Δt = 0.33 s (supplementary method subsection 3). For each polymer configuration, we generated single particle trajectories (SPTs) and computed Lc, kc, D, and alpha (blue traces, Figure 5), which we compared to the experimental data (orange dots, Figure 5). This model was in good agreement with the values of Lc and alpha obtained experimentally, even though only the median distances were used for parameter inference. The agreement was also fair for the spring of constraint Kc. However, we could not recover D.

**Figure 5:**
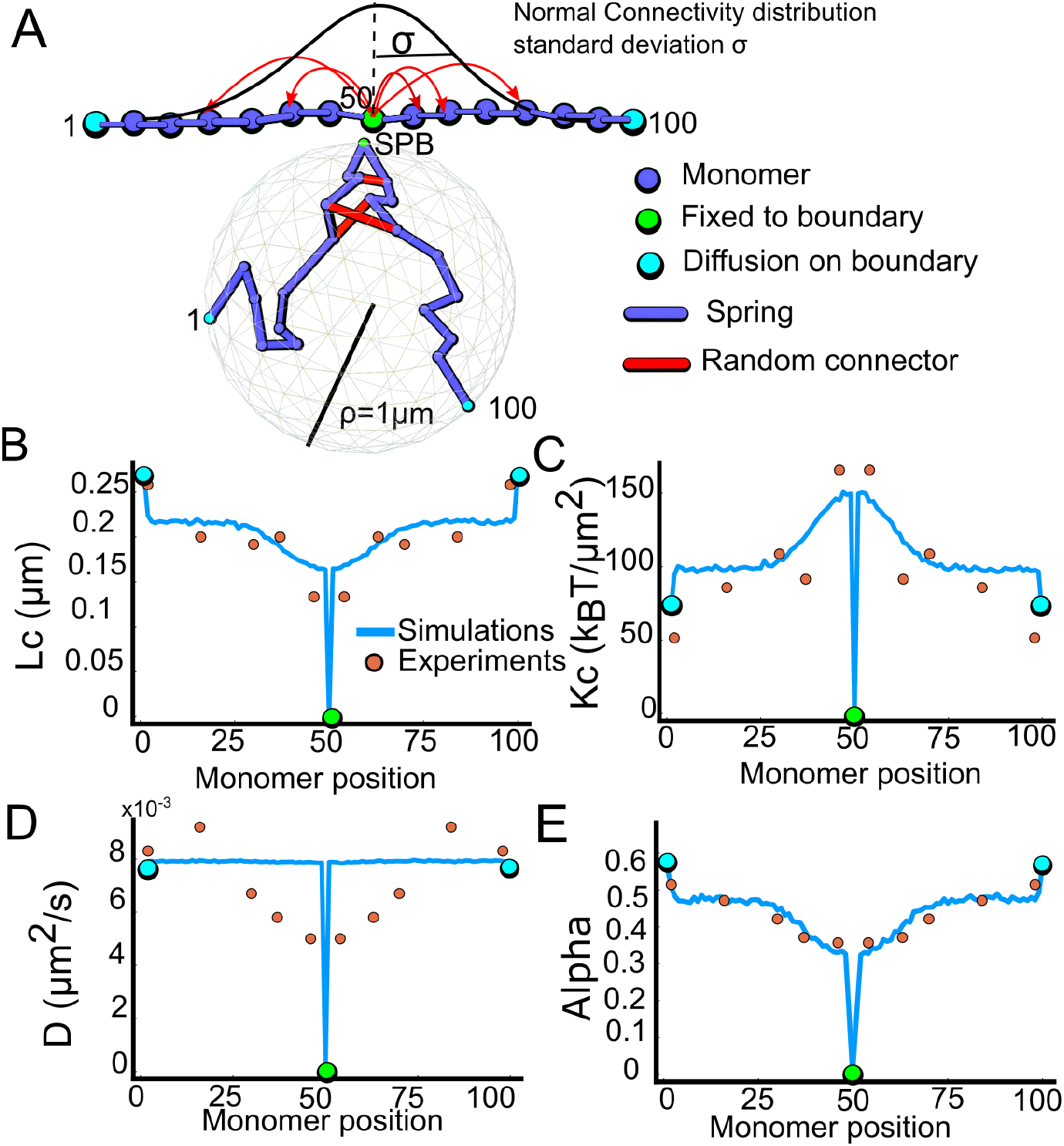
A randomly cross-linked polymer with Gaussian connectivity distribution around the SPB recapitulates chromatin dynamics. A) The chromatin polymer is modelled as a randomly cross-linked polymer with Gaussian connectivity distribution around the SPB. The RCL-polymer is located inside a reflecting spherical domain of radius *ρ*= 1 µm, containing *N* = 100 monomers (purple spheres) connected randomly by connectors (red) around the SPB (monomer 50, green), which is attached at the immobile boundary. *Nc* = 60 connections between monomer pairs, randomly chosen from a normal connectivity distribution with standard-deviation *σ* = 9 (top) centred at the SPB were added. End monomers (cyan, monomers 1 and 100) are freely diffusing on the boundary of the nucleus. B-E) Fluctuation of the indicated parameters obtained with simulations of the polymer described in panel A (blue) for a total of 100 s at time step ∆t=0,03s versus experimental measurements (orange circles): B) Length of confinement Lc, C) Effective spring constant Kc, D) Diffusion coefficient D, E) Anomalous exponent α.

Thus, a modified Randomly Cross-Linked polymer^50^ anchored at a fixed mid-point, with the extremities diffusing on the boundary of a confining sphere or moving freely within the sphere, and Normally distributed connectors from the fixed tether is sufficient to accurately predict the entire statistical distribution observed for Lc, Kc and alpha in vivo. This suggests that physical forces between chromosomes near the SPB constrain motion of whole chromosomes (Figure 5). These tethering forces could be mediated by condensin or cohesin complexes which are able to bridge DNA molecules and are enriched around centromeres in *S. cerevisiae*^51–55^. In addition, the present model suggests that tethering forces decrease rapidly with the distance from the SPB to the telomere. These forces restrict the chromatin motion near the SPB, but were insufficient to further constrain the monomer diffusion rate compared to the experimental results. This difference in the diffusion coefficient might be caused by other mechanisms, such as crowding. Thus, near the SPB, chromosomes seem to be cross-linked over a distance of hundreds of nanometers, impacting large-scale movements.

### Free chromosome extremities exhibit higher mobility

To gain insights into the high mobility of telomeres, we constructed a strain that allowed the induction of a double strand break (DSB) in the middle of the chromosome III arm and analysed whether or not the DSB ends behaved in a similar manner. We followed the ends using two fluorescent tags inserted on either side of the DSB (Figure 6A). Following DSB induction in WT cells the ends do not dissociate, preserving chromosome integrity^56,57^. In yeast, MRX^MRN^ is involved in bridging DSB ends, and cells deficient for *MRE11* exhibit DSB end separation^57,58^. Consistently, we observed DSB end separation in 34% of the *mre11∆* cells (Figure 6A-B). We measured the mobility parameters of separated DSB ends in *mre11∆* cells. As a control, we also imaged *mre11∆* cells in which DSB ends remain together (Figure 6B). Separated ends exhibited a greater spread of trajectories than tethered ends (Figure 6C). As *mre11∆* cells are deficient in non-homologous end joining (NHEJ), which is the main repair pathway in G1 cells, we used *dnl4∆* cells as a control, which are deficient for the NHEJ specific ligase 4, and proficient for DSB ends tethering. The induction of a persistent DSB in *dnl4∆* G1 cells did not significantly affect the spatial constraint, Lc, nor the tethering forces, Kc, acting on the cut site (Figure 6D-E). However, upon cut induction, we observed a significant increase in the diffusion coefficient, D. The nature of the diffusion was also affected as seen in a significant decrease in alpha upon DSB induction in dnl4∆ cells, which is unlikely to be caused by the absence of NHEJ repair in *dnl4∆* mutants as we observed the same decrease in WT cells (Figure S4). Rather, the decrease in alpha could reflect a local change in the chromatin structure. Simulation using the ß-polymer model, where alpha values below 0.5 are associated with compaction of the polymer, has been used to show that polymer decompaction results in increased alpha^46^. Conversely, the decreased alpha values we observed suggest compaction of the chromatin surrounding the DSB in G1 cells.

**Figure 6:**
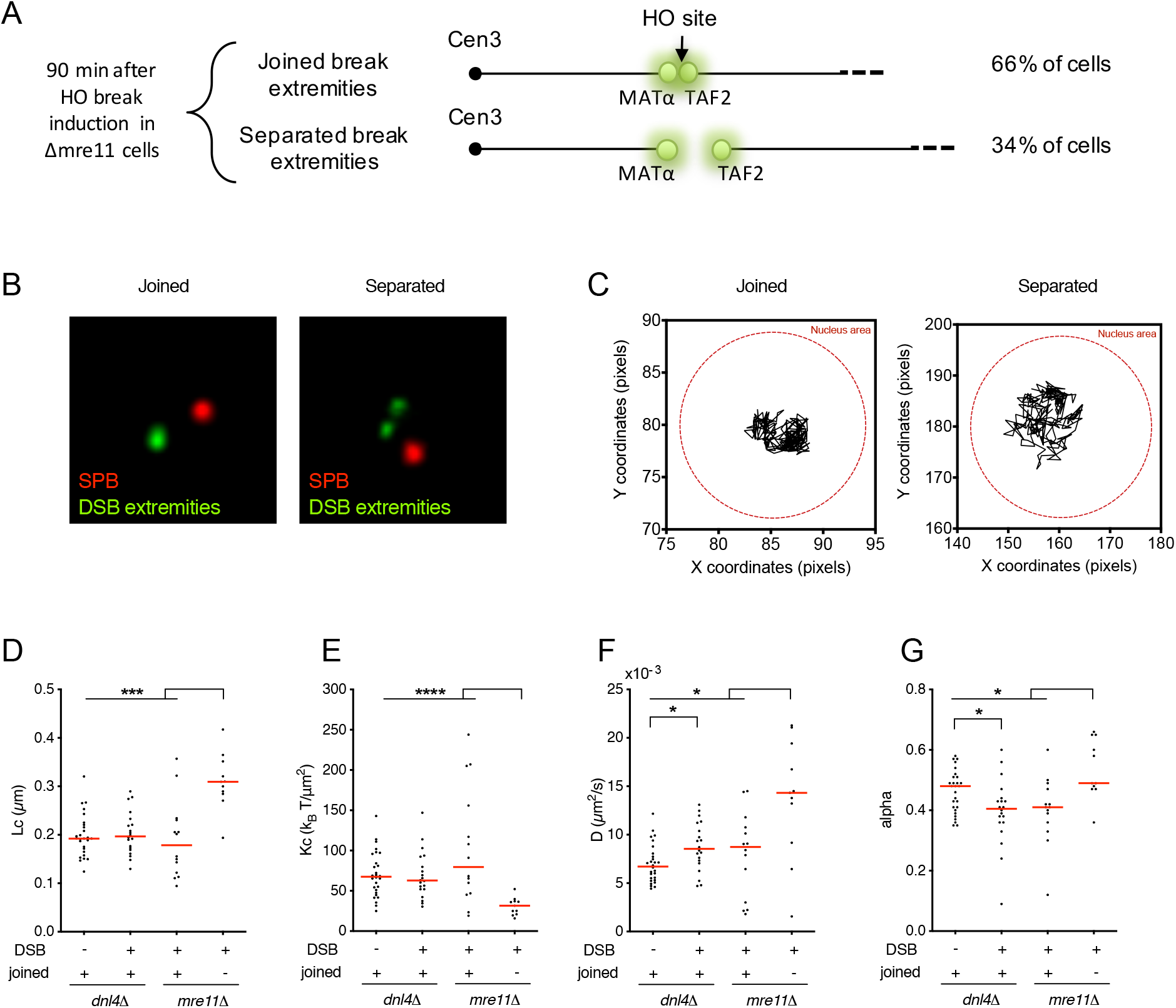
Free chromosome ends exhibit higher mobility. A) System used to follow the two ends of a single double strand break. A DSB is induced at the MAT locus following the galactose-driven expression of the HO endonuclease. *TetO* repeats are inserted at 3.2 kb from the *MAT* locus on the centromere proximal side and *LacO* repeats are inserted 5.2 kb from the *HOcs* site on the telomeric side. After 90 min in galactose the two spots are separated in 34% of the cells in a *∆mre11* strain. B) Micrograph showing a cell with joined DSB ends (one green spot, left panel) and a cell with separated DSB extremities (two green spots, right panel). The SPB is stained in red by the Spc42-mCherry protein. C) Typical trajectories observed for joined or separated extremities. D-G) Distribution of the extracted parameters of the indicated loci and in the growth conditions mentioned. Red bar, distribution mean. Black stars indicate statistical differences (* = p<0,05; ** = p<0,01; *** = p<0,005; **** = p<0,001).

Overall, the DSB dynamics we observed in G1 cells differ from previous reports, which showed that DSB induction increases both the local space explored by the locus (Lc), the diffusion coefficient (D) and the anomalous exponent (alpha)^41,46,59,60^. These differences could arise from differences in the cell cycle, as most of the previous studies analysed DSB motion in S-G2 phase cells that accumulate Rad52 foci, whereas we imaged only G1 phase cells. Thus, DSB induction seem to result in more subtle changes in G1 than in S phase, which affect only the nature of the diffusion and the effective diffusion rate.

Upon break induction, the motion of the tethered DSB ends in *mre11∆* cells was similar to the motion observed in dnl4∆ cells, with undistinguishable Lc, Kc, D and alpha. However, the separated ends exhibited uncoordinated movements and a significant increase in mobility compared to tethered ends (Figure 6C): Lc was increased and Kc concomitantly decreased (Figure 6D-E). The substantial decrease in Kc suggests that the tethering constant also captures chromatin chain constraints, which could represent up to one third of the forces acting on a particular locus. The diffusion coefficient was also substantially increased, from D^joined^ = 7.9 ± 4.3 10^−3^ µm2/s to D^separated^ = 13.8 ± 6.1 10^−3^µm2/s, indicating that the free ends not only explored a larger volume but did so more rapidly (Figure 6F). The anomalous exponent, alpha, increased slightly but significantly for the joined versus separated DSB ends (Figure 6G) and was comparable to the alpha value observed for telomeres (Figure 1H) and for a polymer end moving freely in the nucleus (Figure 4E). Thus, removing the tethering force between the two DSB ends in G1 cells releases some of the constraints imposed by neighbouring DNA and lead to faster motion in a larger volume.

The dynamic parameters Lc, Kc and alpha observed for free DSB ends compared well with those observed for freely diffusing telomeres (TEL6Ri; Figure 2D-G). However, the diffusion coefficient was significantly higher for free DSB ends, with D= 13 ± 6.12 10^−3^ µm^2^/s, compared to TEL6Ri, with D= 9 ± 1.77 10^−3^ µm^2^/s, suggesting that telomeres are subjected to greater resistance to motion. This may reflect that intranuclear telomeres show a lower degree of compaction and are thus more sensitive to nucleoplasm viscosity.

## Discussion

We have analysed the dynamics of yeast single loci in the yeast genome by fast live imaging, characterizing their mobility using four biophysical parameters: the length of confinement of the trajectory (Lc); the tethering constant (Kc); the diffusion coefficient (D); and the anomalous diffusion exponent, alpha^46^. While our results recapitulate previous data at a global level^2,16,39,40^, we observed quantitative differences at a local level that may be accounted for by changes in several experimental parameters. First, we labelled loci using a low affinity LacO which is less likely to perturb chromatin structure^47^. Second, we confined our analysis to G1 cells and only analysed movies with low parasitic rotations or oscillations of the nucleus. Despite this precaution, we observed large cell-to-cell variations for each parameter. This suggests that chromatin motion is influenced by stochastic events, such as changes in the local chromatin folding and environment, in transcription events or interactions with nuclear structures, which vary from cell to cell independently of cell cycle stage.

Chromatin interactions with nuclear structures, such as centromere anchoring to the SPB in yeast, have been shown to constraint the space explored by loci near the tethering point^2^. Our data extend this observation, showing that loci up to 33 kb away from the centromere have a low alpha and a low diffusion coefficient compared to more distant loci. We also observed that telomeres moving on the NE are no more confined than intrachromosomal loci. Consistently, after detaching a telomere *TEL6R* from the membrane, its dynamics increase, while adding an intermediate anchoring point (1.8 kb away from the fluorescent tag on *TEL6R*) confined the telomere motion and reduce the dynamics. Thus, chromatin movement is constrained by tethering to the NE, although to a less extent than tethering to centromeres, consistent with the transient nature of telomere-NE interactions. In addition to their interaction with the NE, telomeres interact together in Sir3 mediated clusters from which they dissociate/associate on a time scale of tenth of seconds^61–63^. The mechanism of telomere dissociation from clusters remains unclear, but this transient weak interaction is also described by the tethering constant Kc which for peripheral telomeres is half that of centromeres. Adding a second interaction point with the NPC decreases mobility and results in doubling the tethering constraint Kc. This doubling suggests that telomere-NE and gene-NPC interactions occur with equivalent forces. Constrained dynamics of loci close to the NE has also been observed in mammalian cells^61,64^. The estimation of the tethering constraint as defined here could yield insights into the nature of the chromosome-NE interaction.

Simplified polymer models have been used to recapitulate the nuclear organisation and dynamics of the yeast genome^2,34,42,65–67^. Using an elementary Rouse polymer model for chromosomes confined into the nucleus, with fixed tethering for the centromeres and telomeres, was able to capture the tethering constraint of the centromeres^2,42,44^ but was largely incapable of recapitulating the variation in Lc, Kc, D and alpha along the chromosome. We therefore constructed a more sophisticated polymer model, adding cross-linkers normally distributed around the centromeric region, which was sufficient to accurately three of the four parameters. Candidate for these cross-linkers include structural maintenance of chromosomes (SMC) family proteins, which ring-shape allows to encircle two DNA strands and form loops^68^. Among SMCs, the cohesin complex is loaded onto the core centromeric region and spreads into the surrounding pericentromere but this only occurs in late G1^51–53,69^. The condensin complex is similarly enriched around the centromere including in G1 making it a good candidate to test^54,55^.

Despite the addition of cross-linkers, our model did not recapitulate the observed variation in the diffusion coefficient, D. D is sensitive to crowding, which can could occur at different spatial scales from tens to hundreds of nanometres, yet, attempts to introduce crowding into our model failed to recapitulate the diffusion rate along the chromosome, possibly because D is also very sensitive to the local environement. We observed that the coefficient of diffusion of the trnascriptionaly activated *HXK1* gene decreases independently of its interaction with the nuclear periphery. This is reminiscent of the confined motion of mRNA-producing genes observed in human cells^70^ and agrees with the increased chromatin dynamics observed upon transcription arrest^71^. Transcriptional activation and the action of nucleosome remodelers have in other instances been shown to increase chromatin diffusion coefficient in yeast^72^. Thus, chromatin diffusion coefficient seems sensitive to local perturbations that can vary according to the position on the chromosome and also from cell-to-cell depending on the position of the locus relative to nuclear structures or to other genomic elements, making it difficult to model. The telomere motion we observe here is not consistent with fixed tethers at the nuclear membrane but is best described by the motion modelled for the cross-linked polymer, where the ends can move freely within the nucleus or slide unhindered across the NE. Our model also revealed that the polymer ends exhibit an overall higher mobility than internal monomers suggesting that the high mobility observed for telomeres may be an intrinsic property of chromosome ends. Cas9-GFP or TALE-GFP marked telomeres are seen to have high mobility within the nuclear interior of human cells^73^, and we observed that chromosome ends produced by the induction of a DSB in the middle of a chromosome arm also exhibit increased mobility, similar to that of freely moving telomeres.

Altogether, these results suggest that the observed telomere dynamics are an intrinsic propensity of chromosome ends to be highly mobile, which is modulated by transient interactions with proteinous compartments and fixed nuclear components such as the NE.

We report here that induction of a single DSB in G1 cells does not affect the mobility parameters as previously observed in S phase cells. Indeed,, Induction of a single DSB in S phase cells^41,59,60^ results in the DSB ends showing an increase in: the space they explored, theit velocity, as measured by the effective diffusion coefficient, and the anomalous exponent, alpha, which based on polymer modelling, was suggested to result from a local chromatin decompaction^46^. In contrast, our data show that DSBs induced in G1 cells do not increase DSB end exploration but instead lead to a significant decrease in alpha, indicating G1 DSBs explore space in a more recurrent fashion, suggesting that local compaction of the chromatin fibre occurs during this phase of the cell cycle, reminiscent of the transient and local chromatin compaction seen at DSB in mammalian cells^74–78^. Since NHEJ is the preferred repair pathway in G1, we speculate that a local compaction could prevent large scale movements of the damaged domain so that the DSB ends remain in close proximity, increasing the likelihood of end joining. Consistently, after separation, both the space explored by DSB ends and their diffusion rate increase substantially.

Loss of mobility following DSB induction has been observed in cells prior to resection, the processing step producing the ssDNA required to search for a homologous sequence and to prime synthesis for homologous recombination^79^. This mobility loss is consistent with the constrained dynamics we observed in G1, where resection is repressed. Resection forms the ssDNA, which is coated by the Rad51 recombinase. At this stage, several studies have shown that DSB mobility increases, which may result from stiffening of the ssDNA by Rad51 binding^80^, or stiffening of the surrounding chromatin^81^ or a local chromatin decompaction^46,82^. Damaged chromatin dynamics seem to evolve as part of a multistep process that is regulated across the cell cycle, consistent with the preferred pathways in the two cell cycle phases: in S phase, increased mobility could favour homology searching and HR, whereas in G1 the two ends should stay together to allow rejoining through NHEJ.

To conclude, we demonstrated here that, in yeast, both telomere and DSB free ends exhibit high mobility, likely related to their position at the ends of the chromosomes, a property that could be evolutionary conserved since damage foci and telomeres also exhibit enhanced mobility compared to interstitial loci in human cells^73,83^.

## Material and Methods

### Yeast and growth conditions

Yeast strains used in this study are all derivatives of the JKM179 strain ^84^ which is *MATα ade1 leu2-3, leu2-112 lys5 trp1::hisG ura3-52* and are listed in Table S1. Strains were obtained through insertion of both a Lac operator array (256 lacOp repeats) at the indicated chromosomal location, a Nup49-mCherry fusion and a non-tetramerizing low affinity DNA binding lac repressor-GFP fusion (GFP-LacI**) under the *HIS3* promoter ^47^. The affinity of the lacI** allele is lowered by the insertion of three glycines replacing Gln^60^ in the hinge region, downstream of the DNA binding domain ^47,85^. In addition, a R197A mutation that lowers lacI binding to inducer sugars ^86^ was introduced to allow imaging in galactose containing medium. The *lacO* or *TetO* arrays were integrated in chromosome VII at coordinate 484500 (12.5 kb from *CEN7*), in chromosome III at position 148110 (0.5 kb from *CWH43*), 197344 (3.2 kb upstream of HOcs at *MAT*) and 206118 (0.73 kb from TAF2, 5.2 kb downstream of HOcs at *MAT*), in chromosome VI at coordinate 198838 (0.8 kb from *PES4*; ^14^) and at coordinate 256905 (13 kb from *TEL6R*; ^40^).

To serve as a static reference point in the nucleus, the Spc42 protein was fused to yEGFP. Gene deletions were obtained through a PCR-mediated transformation as described in (Longtine et al., 1998). All insertions or deletions were verified by PCR and phenotypic assays. Cells were grown to logarithmic phase at 30°C in complete synthetic media with 2% glucose or 2% galactose. For DSB induction, cells were grown in 2mL in rich medium (yeast extract– peptone–dextrose, YPD) overnight. Cultures were then diluted in rich medium containing 2% lactate, 3% glycerol, 0.05% glucose and grown to OD_600_=0.3-0.8. HO was induced by addition of galactose to a final concentration of 2% for 2h prior imaging. The efficiency of HO endonuclease cleavage at *MAT* was determined by quantitative PCR as described in ^87^.

### Microscopy acquisition and analysis

Microscopy Images were captured with a ×100 magnification oil-immersion objective (1.46 numerical aperture) on a Leica DMI 6000B microscope (Leica Microsystems) equipped with a piezoelectric translator (PIFOC, Physik Instrumente), a ORCA-Flash 4.0 camera (Hamamatsu), an illumination system with leds (Lumencore) and rapid imaging software (Metamorph). Wavelengths of the leds used are 475nm (for GFP, 205mW), and/or 575nm (for mCherry, 300MW). Two-minute movies were acquired with stacks of 10 optical slices separated by 250nm every 338ms. Each slice was exposed for 30ms for a total of 330ms per stack. All microscopy was done in microfluidic plates (Y04C plates, ONIX platform, CellASIC) in a temperature-controlled environment set to 25°C. The raw images were deconvolved using the Autoquant software. The movies were then tracked using ImageJ ^88^ with the Mosaic macro ^89^ to produce 3D+t trajectories. Further processing and analysis of the movies was done using Matlab.

LacI-GFP position relative to the nuclear membrane was determined with a through-focus stack of twenty 0.2 µm steps and was measure using the ImageJ plug-in software PointPicker. To determine zone enrichment, we applied a proportional analysis comparing zone 1 with a random distribution or the enrichment in zone 1 of two different strains with a confidence limit of 95%.

### Extraction of parameters, modelling and simulations with polymer models

The biophysical parameters Lc, Kc, D and alpha extracted from the trajectories obtained from the image stacks are described in the supplementary methods (section 2) see also ^46^.

The polymer models used here are described in the supplementary methods (section 1 and 3). Briefly, we used Rouse polymer model which is a collection of monomers moving with a random Brownian motion coupled to a spring force originating from the nearest neighbours. We modified Rouse model by adding connectors (RCL polymer model as described in ^50^). We improved here this model by adding connectivity distributed according to a Gaussian distribution (supplementary methods section 3).

### Statistical analysis

Significance between distribution of distances from the nuclear membrane was tested by a proportional analysis comparing the percentage of distances below 200nm. Statistical significance was determined by using a 95% confidence interval.

Significance between a, Kc, Lc, and D values was tested using the non-parametric Kolmogorov-Smirnov (KS) test. p values are either indicated above the figure panels or by asterisks, where *p % 0.05, **p % 0.01, ***p% 0.001, respectively. All p values are given in supplementary Table S2 to S6.

## Supporting information

Toulouze_supplementary_material

## Author contributions

M. T. and K. D. planned the experiments. M.T. performed most experiments. M. T., K. D., and A. A. analysed the imaging data. A. A. and D. H. carried out the theoretical development of the analysis pipeline. A. A. extracted biophysical parameters from imaging data. A. A., O. S. and D. H. carried out polymer modelling. A. A. and O. S. carried out polymer simulations. K. D. and D. H. wrote the paper with inputs from all the authors.

## Acknowledgments

We thank S. Marcand for critically reading this manuscript and all the Dubrana and Holcmann laboratory’s member for fruitful discussions. K.D. thanks the European Research Council under the European Community’s Seventh Framework Program (FP7/2007 2013/European Research Council grant agreement 281287), DRF-Impulsion (4D-DSB-DIC) and the Radiobiology program of the CEA Segment 4 for support. D. Holcman is supported by a team funding from the Fondation pour la Recherche Médicale (FRM).

