## Supplementary material for "Telomeric chromosome ends are highly mobile and behave like free double-strand DNA breaks": Toulouze_supplementary_material

#### Inventory

##### 1- Supplemental Tables

*Table S1: S. cerevisiae strains used in this paper*

A list of the yeast strains, index number and original citation.

*Table S2: p values relative to figure 1*

Statistical analysis of the data presented in Figure 1 using a non-parametric Kolmogorov-Smirnov (KS) test.

*Table S3: p values relative to figure 2*

Statistical analysis of the data presented in Figure 2 using a non-parametric Kolmogorov-Smirnov (KS) test.

*Table S4: p values relative to figure 3 D-G*

Statistical analysis of the data presented in Figure 3 using a non-parametric Kolmogorov-Smirnov (KS) test.

*Table S5: p values relative to figure 3 I-P*

Statistical analysis of the data presented in Figure 3 using a non-parametric Kolmogorov-Smirnov (KS) test.

*Table S6: p values relative to figure 6*

Statistical analysis of the data presented in Figure 6 using a non-parametric Kolmogorov-Smirnov (KS) test.

##### 2- Supplementary Figures

*Figure S1: relates to Figure 1*

*Figure S2: relates to Figure 1*

*Figure S3: relates to Figure 1 and S2*

*Figure S4: relates to Figure 2*

*Figure S5: relates to Figure 6*

##### 3- Supplementary Methods

### 1. Toulouze et al. Supplemental Tables

#### Table S1: *S. cerevisiae* strains used in this paper:

| Strain | Genotype | Reference |
|---|---|---|
| JKM179 | *MATα hml::ADE1 hmr::ADE1 ade3::pGal-HO ade1 leu2-3,112 lys5 trp1::hisG ura3-52* | Lee et al. 1998 |
| JKM139 | *MATα hml::ADE1 hmr::ADE1 ade3::pGal-HO ade1 leu2-3,112 lys5 trp1::hisG ura3-52* | Lee et al. 1998 |
| yKD385 | JKM179 *MATα::lacOp-TRP1 leu2-3::GFP-lacI-LEU2 NUP49::NUP49-mCherry-URA3 SPC42-yEGFP-HPH* | This study (Figure S3) |
| yKD388 | JKM179 *matα-inc::lacOp-TRP1 leu2-3::GFP-lacI-LEU2 NUP49::NUP49-mCherry-URA3 SPC42-yEGFP-HPH* | This study (Figure S3) |
| yKD539 | JKM179 *MAT::lacOpFX-TRP1 leu2-3::GFP-lacI**R-LEU2 NUP49::NUP49-mCherry-URA3 SPC42-yEGFP-HPH dnl4::KanMx* | This study (Figure 5) |
| yKD667 | JKM179 *matα-inc::lacOpFX-TRP1 leu2-3::GFP-lacI**R-LEU2 NUP49::NUP49-mCherry-URA3 SPC42-yEGFP-HPH* | This study (Figure 1, 4) |
| yKD724 | JKM179 TEL6R*::lacOp-TRP1 leu2-3::GFP-lacI**R-LEU2 NUP49::NUP49-mCherry-URA3 SPC42-yEGFP-HPH* | This study (Figure 1, 2, 3, 4, S2) |
| yKD732 | JKM179 *matα-inc::lacOpFX-TRP1 leu2-3::GFP-lacI**R-LEU2 NUP49::NUP49-mCherry-URA3 SPC42-yEGFP-HPH dnl4::KanMx* | This study (Figure 5) |
| yKD750 | JKM179 *matα-inc TEL6R::lacOp-TRP1 leu2-3::GFP-lacI**R-LEU2 NUP49::NUP49-mCherry-URA3 SPC42-yEGFP-HPH* | This study (Figure 3) |
| yKD836 | JKM179 *TEL6R::lacOp-TRP1 leu2-3::GFP-lacI**R-LEU2 NUP49::NUP49-mCherry-URA3 SPC42-yEGFP-HPH ku80::KanMx* | This study (Figure 3) |
| yKD890 | JKM179 *matα-inc TEL6R::lacOp-TRP1 leu2-3::GFP-lacI**R-LEU2 NUP49::NUP49-mCherry-URA3 SPC42-yEGFP-HPH hxk1::KanMx* | This study (Figure 3) |
| yKD985 | JKM139 *0.5kbCWH43::lacOpFX-TRP1 ura3-52::lacI-mCherry-URA3 SPC42-yEGFP-HPH* | This study (Figure 1, 4) |
| yKD1050 | JKM179 *PES4::lacOp-TRP1 leu2-3::GFP-lacI**R-LEU2 NUP49::NUP49-mCherry-URA3 SPC42-yEGFP-HPH* | This study (Figure 1, 2, 3, 4, S2) |
| yKD1112 | W303 *12kbCEN7::lacOp-TRP1 ura3-1::GFP-lacI**R-URA3 SPC42-mCherry-HPH 2pGal-CEN6* | This study (Figure 1, 4) |
| yKD1113 | JKM139 *TAF2-lacOpFx-TRP1 MAT-TetO-LEU2 ura3-1::GFP-lacI**R-URA3 leu2::TetR-GFP-LEU2 Spc42-mCherry::KanMX mre11::HPH* | This study (Figure 5) |

#### Table S2: p values relative to figure 1

| Lc | CEN7 (12kb) | CWH43 (33kb) | PES4 (50kb) | MAT (86kb) | TEL6R (115kb) |
|---|---|---|---|---|---|
| CEN7 (12kb) | | 0,0027 | 0,0075 | < 0,0001 | < 0,0001 |
| CWH43 (33kb) | | | 0,3958 | 0,7805 | 0,0286 |
| PES4 (50kb) | | | | 0,3957 | 0,0033 |
| MAT (86kb) | | | | | 0,0052 |
| TEL6R (115kb) | | | | | |

| Kc | CEN7 (12kb) | CWH43 (33kb) | PES4 (50kb) | MAT (86kb) | TEL6R (115kb) |
|---|---|---|---|---|---|
| CEN7 (12kb) | | 0,0167 | 0,0336 | 0,0007 | < 0,0001 |
| CWH43 (33kb) | | | 0,8402 | 0,6102 | 0,0072 |
| PES4 (50kb) | | | | 0,2324 | 0,0022 |
| MAT (86kb) | | | | | 0,0049 |
| TEL6R (115kb) | | | | | |

| D | CEN7 (12kb) | CWH43 (33kb) | PES4 (50kb) | MAT (86kb) | TEL6R (115kb) |
|---|---|---|---|---|---|
| CEN7 (12kb) | | 0,2283 | 0,0292 | < 0,0001 | < 0,0001 |
| CWH43 (33kb) | | | 0,3001 | 0,0004 | 0,0009 |
| PES4 (50kb) | | | | 0,007 | 0,0464 |
| MAT (86kb) | | | | | 0,3994 |
| TEL6R (115kb) | | | | | |

| alpha | CEN7 (12kb) | CWH43 (33kb) | PES4 (50kb) | MAT (86kb) | TEL6R (115kb) |
|---|---|---|---|---|---|
| CEN7 (12kb) | | 0,934 | 0,0002 | 0,0175 | < 0,0001 |
| CWH43 (33kb) | | | 0,0031 | 0,134 | 0,0001 |
| PES4 (50kb) | | | | 0,2841 | 0,1295 |
| MAT (86kb) | | | | | 0,0194 |
| TEL6R (115kb) | | | | | |

#### Table S3: p values relative to figure 2

| Lc | PES4 | TEL6R | TEL6Rp | TEL6Ri |
|---|---|---|---|---|
| PES4 | | 0,0033 | 0,2155 | 0,0011 |
| TEL6R | | | 0,1737 | 0,3191 |
| TEL6Rp | | | | 0,0403 |
| TEL6Ri | | | | |

| Kc | PES4 | TEL6R | TEL6Rp | TEL6Ri |
|---|---|---|---|---|
| PES4 | | 0,0022 | 0,1889 | 0,0007 |
| TEL6R | | | 0,1737 | 0,3191 |
| TEL6Rp | | | | 0,0408 |
| TEL6Ri | | | | |

| D | PES4 | TEL6R | TEL6Rp | TEL6Ri |
|---|---|---|---|---|
| PES4 | | 0,3994 | 0,085 | 0,9815 |
| TEL6R | | | 0,0725 | 0,1907 |
| TEL6Rp | | | | 0,006 |
| TEL6Ri | | | | |

| alpha | PES4 | TEL6R | TEL6Rp | TEL6Ri |
|---|---|---|---|---|
| PES4 | | 0,1295 | 0,8068 | 0,0495 |
| TEL6R | | | 0,2896 | 0,4364 |
| TEL6Rp | | | | 0,1114 |
| TEL6Ri | | | | |

#### Table S4: p values relative to figure 3 D-G

| Lc | TEL6R ku80 | TEL6R p | TEL6R i |
|---|---|---|---|
| TEL6R ku80 | | 0,0184 | 0,6328 |
| TEL6R p | | | 0,0403 |
| TEL6R i | | | |

| Kc | TEL6R ku80 | TEL6R p | TEL6R i |
|---|---|---|---|
| TEL6R ku80 | | 0,0454 | 0,5915 |
| TEL6R p | | | 0,0408 |
| TEL6R i | | | |

| D | TEL6R ku80 | TEL6R p | TEL6R i |
|---|---|---|---|
| TEL6R ku80 | | 0,0983 | 0,9246 |
| TEL6R p | | | 0,006 |
| TEL6R i | | | |

| alpha | TEL6R ku80 | TEL6R p | TEL6R i |
|---|---|---|---|
| TEL6R ku80 | | 0,4273 | 0,6034 |
| TEL6R p | | | 0,1114 |
| TEL6R i | | | |

#### Table S5 : p values relative to figure 3 I-P

| Lc | TEL6R GLU | TEL6Rp GLU | TEL6Ri GLU | TEL6R GAL | TEL6R hxk1 GAL | TEL6Rp GAL | TEL6Ri GAL |
|---|---|---|---|---|---|---|---|
| TEL6R GLU | | 0,1737 | 0,3191 | 0,0033 | 0,6299 | < 0,0001 | 0,3191 |
| TEL6Rp GLU | | | 0,0403 | 0,4691 | 0,0774 | 0,0392 | 0,2042 |
| TEL6Ri GLU | | | | 0,0003 | 0,5712 | 0,0008 | 0,1418 |
| TEL6R GAL | | | | | 0,0007 | 0,1443 | 0,0508 |
| TEL6R hxk1 GAL | | | | | | < 0,0001 | 0,2869 |
| TEL6Rp GAL | | | | | | | 0,0025 |
| TEL6Ri GAL | | | | | | | |

| Kc | TEL6R GLU | TEL6Rp GLU | TEL6Ri GLU | TEL6R GAL | TEL6R hxk1 GAL | TEL6Rp GAL | TEL6Ri GAL |
|---|---|---|---|---|---|---|---|
| TEL6R GLU | | 0,1737 | 0,3191 | 0,0084 | 0,8947 | < 0,0001 | 0,8916 |
| TEL6Rp GLU | | | 0,0408 | 0,5495 | 0,1723 | 0,0271 | 0,1652 |
| TEL6Ri GLU | | | | 0,0011 | 0,232 | < 0,0001 | 0,2302 |
| TEL6R GAL | | | | | 0,0149 | 0,1097 | 0,032 |
| TEL6R hxk1 GAL | | | | | | < 0,0001 | 0,9789 |
| TEL6Rp GAL | | | | | | | 0,0008 |
| TEL6Ri GAL | | | | | | | |

| D | TEL6R GLU | TEL6Rp GLU | TEL6Ri GLU | TEL6R GAL | TEL6R hxk1 GAL | TEL6Rp GAL | TEL6Ri GAL |
|---|---|---|---|---|---|---|---|
| TEL6R GLU | | 0,0725 | 0,1907 | 0,0008 | 0,7069 | 0,0006 | 0,1907 |
| TEL6Rp GLU | | | 0,006 | 0,1213 | 0,0471 | 0,0606 | 0,5569 |
| TEL6Ri GLU | | | | 0,0004 | 0,3844 | 0,0003 | 0,0355 |
| TEL6R GAL | | | | | 0,0004 | 0,469 | 0,3364 |
| TEL6R hxk1 GAL | | | | | | 0,0008 | 0,1097 |
| TEL6Rp GAL | | | | | | | 0,1469 |
| TEL6Ri GAL | | | | | | | |

| alpha | TEL6R GLU | TEL6Rp GLU | TEL6Ri GLU | TEL6R GAL | TEL6R hxk1 GAL | TEL6Rp GAL | TEL6Ri GAL |
|---|---|---|---|---|---|---|---|
| TEL6R GLU | | 0,2896 | 0,4364 | 0,0185 | 0,2609 | 0,0142 | 0,1237 |
| TEL6Rp GLU | | | 0,1114 | 0,3989 | 0,9089 | 0,331 | 0,5119 |
| TEL6Ri GLU | | | | 0,0079 | 0,1274 | 0,0048 | 0,0729 |
| TEL6R GAL | | | | | 0,1906 | 0,8264 | 0,7641 |
| TEL6R hxk1 GAL | | | | | | 0,1237 | 0,6178 |
| TEL6Rp GAL | | | | | | | 0,6553 |
| TEL6Ri GAL | | | | | | | |

#### Table S6 : p values relative to figure 6

| Lc | *dnl4Δ* +DSB | *dnl4Δ* -DSB | *mre11Δ* +DSB joined | *mre11Δ* +DSB separated |
|---|---|---|---|---|
| *dnl4Δ* +DSB | | 0,7293 | 0,3182 | < 0,0001 |
| *dnl4Δ* -DSB | | | 0,3938 | < 0,0001 |
| *mre11Δ* +DSB joined | | | | 0,0018 |
| *mre11Δ* +DSB separated | | | | |

| Kc | *dnl4Δ* +DSB | *dnl4Δ* -DSB | *mre11Δ* +DSB joined | *mre11Δ* +DSB separated |
|---|---|---|---|---|
| *dnl4Δ* +DSB | | 0,7997 | 0,2861 | < 0,0001 |
| *dnl4Δ* -DSB | | | 0,3115 | < 0,0001 |
| *mre11Δ* +DSB joined | | | | 0,0007 |
| *mre11Δ* +DSB separated | | | | |

| D | *dnl4Δ* +DSB | *dnl4Δ* -DSB | *mre11Δ* +DSB joined | *mre11Δ* +DSB separated |
|---|---|---|---|---|
| *dnl4Δ* +DSB | | 0,0138 | 0,7061 | 0,0063 |
| *dnl4Δ* -DSB | | | 0,2712 | 0,0005 |
| *mre11Δ* +DSB joined | | | | 0,021 |
| *mre11Δ* +DSB separated | | | | |

| alpha | *dnl4Δ* +DSB | *dnl4Δ* -DSB | *mre11Δ* +DSB joined | *mre11Δ* +DSB separated |
|---|---|---|---|---|
| *dnl4Δ* +DSB | | 0,0223 | 0,8289 | 0,0008 |
| *dnl4Δ* -DSB | | | 0,0677 | 0,0431 |
| *mre11Δ* +DSB joined | | | | 0,0047 |
| *mre11Δ* +DSB separated | | | | |

#### 2. Toulouze et al. Supplementary figures

**Supplementary figure legends:**

**Figure S1:**

A) Representative Image showing the nuclear periphery labelled with Nup49-mCherry, the SPB labelled by SPC42-GFP and a LacO-LacI-GFP tagged locus.

B) acquisition scheme and SPB, spot trajectories projected on a gray scaled imaged of the Nup49-mCherry tagged nuclear periphery.

C-F) Correlation plot of the Lc, Kc, D and alpha parameters with SPB Lc. Red line, distribution median of the Lc, Kc, D and alpha parameters. Grey dotted line, SPB Lc threshold used to sort the movies. Only movies with mean SPB Lc below 200 nm were analysed. Indeed, SPB Lc above 200 nm show high Lc and low Kc suggesting that parasitic nuclear movement are occurring.

**Figure S2:**

A) Scheme of the simulations. The end monomer of a Rouse polymer is attached to an surface oscillating with a sinusoidal motion. The Rouse polymer is located inside a reflecting spherical domain of radius $\rho$ = 1m, containing N = 100 monomers (purple spheres) connected randomly by connectors (red) around the SPB (monomer 50, green), which is attached at the immobile boundary.

B-E) For a fixed amplitude angle $\theta_{\max} = \frac{\pi}{8}$, evolution of the four parameters obtained for each monomer for various frequencies $\omega$ varying in the range in [0 -12] Hz.

F-I) For a fixed frequency $\omega$ = 30Hz, plot of the four parameters for each monomer for an angular amplitude varying in the range of [0,3] μm.

The oscillations of the boundary influence telomere motion, but not loci located at a genomic distance of 10% of the total length of an anchored point. Note however, the large fluctuations between different realizations (Fig. S3)

**Figure S3: Single realization analysis for a Rouse polymer attached to an oscillating nuclear envelop.**

A) Scheme of the simulations. The end monomer (green) is attached to an oscillating surface. The Rouse polymer

is located inside a reflecting spherical domain of radius= 1 µm, containing N = 100 monomers (purple spheres) connected randomly by connectors (red) around the SPB (monomer 50, green), which is attached at the immobile
boundary.

B-E) For a fixed frequency $\omega$ = 8Hz and amplitude angle $\theta_{\max} = \frac{\pi}{8}$, plot for each realization of the four parameters (mean in dashed line): B. Length of confinement Lc, C. Effective spring constant Kc, D. Diffusion coefficient D. E. Anomalous exponent.

**Figure S4:**

A) Distribution of the spot to membrane distance for each *TEL6R* movie analysed.

B) Distribution of the spot to membrane distance for each *PES4* movie analysed.

C-F) Cumulative distribution function for the distance of the spot to the NE along each the movies analysed for *TEL6R* (C), *TEL6R* movies with a mean spot-membrane distance <200nm (D), *TEL6R* movies with a mean spot-membrane distance >200nm (E), for *PES4* movies (F).

**Figure S5:**

A-D) Distribution of the extracted parameters of the MAT locus upon growth in galactose for a strain with an uncleavable locus (-DSB) and for a cleavable locus (+DSB). Red bar, distribution mean. Black stars indicate statistical differences (* = p<0,05; ** = p<0,01; *** = p<0,005; **** = p<0,001).

**Figure S1**

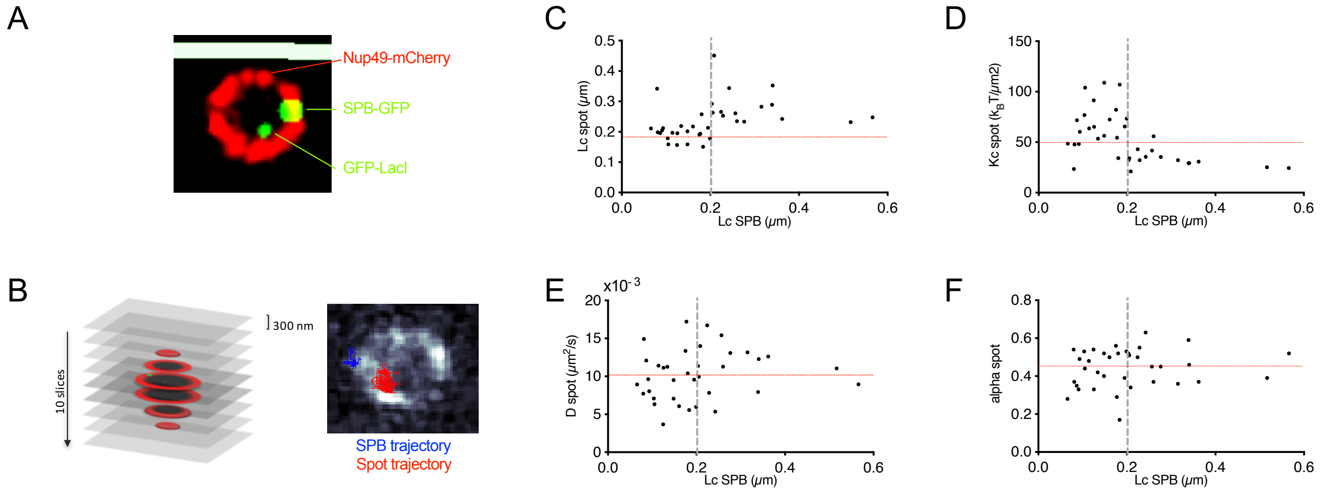
A
Nup49-mCherry
SPB-GFP
GFP-LacI
B
300 nm
10 slices
SPB trajectory
Spot trajectory
C
Lc spot (µm)
Lc SPB (µm)
D
Kc spot (kBT/µm2)
Lc SPB (µm)
E
x10-3
D spot (µm2/s)
Lc SPB (µm)
F
alpha spot
Lc SPB (µm)

**Figure S2**

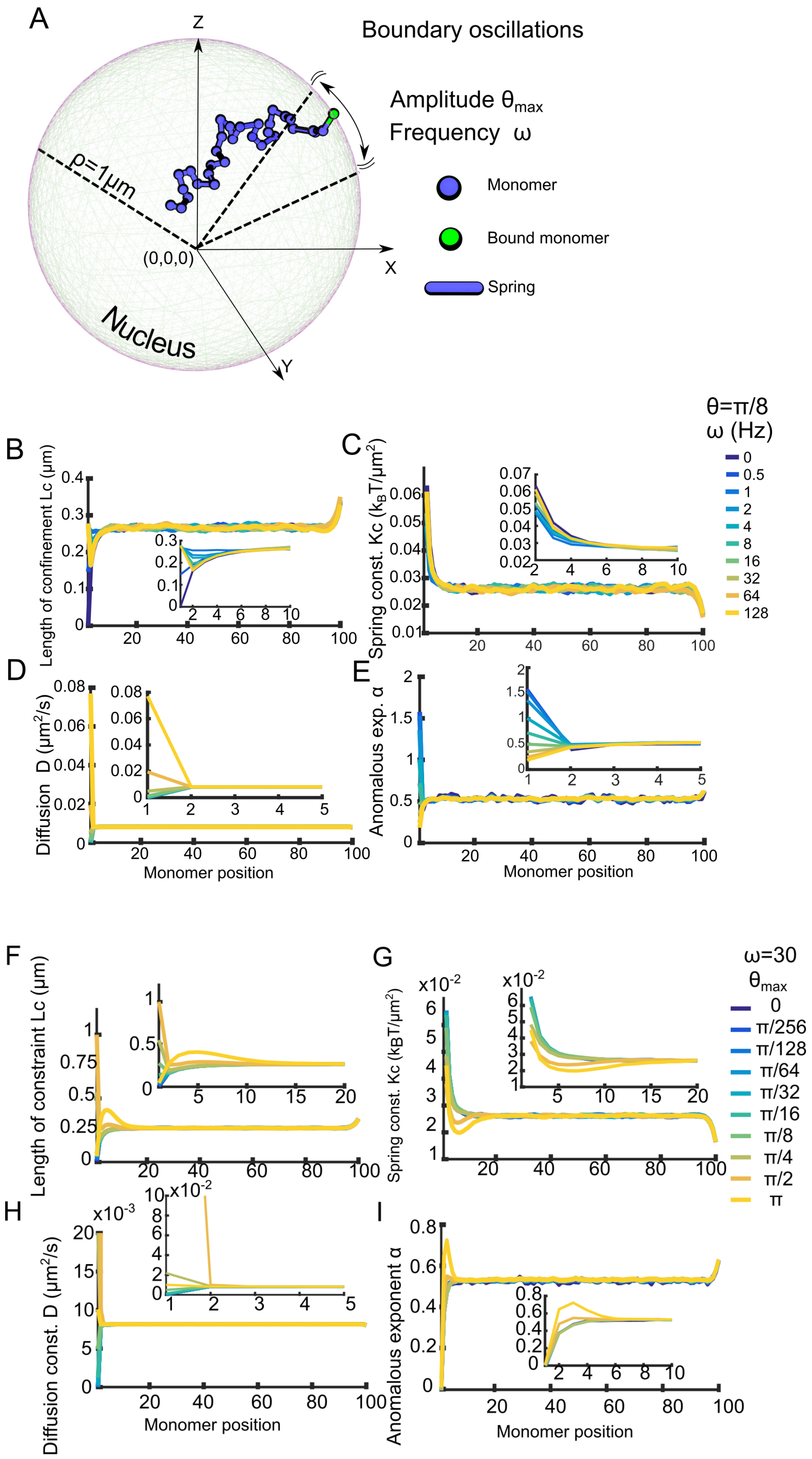
A
Boundary oscillations
Amplitude $\theta_{max}$
Frequency ω
Monomer
Bound monomer
Spring
Z
X
Y
ρ=1μm
(0,0,0)
Nucleus
B
C
θ=π/8
ω (Hz)
0
0.5
1
2
4
8
16
32
64
128
Length of confinement Lc (μm)
Spring const. Kc ($k_BT/\mu m^2$)
D
E
Diffusion D ($\mu m^2/s$)
Anomalous exp. α
Monomer position
F
G
ω=30
$\theta_{max}$
0
π/256
π/128
π/64
π/32
π/16
π/8
π/4
π/2
π
Length of constraint Lc (μm)
Spring const. Kc ($k_BT/\mu m^2$)
H
I
Diffusion const. D ($\mu m^2/s$)
Anomalous exponent α
Monomer position

**Figure S3**

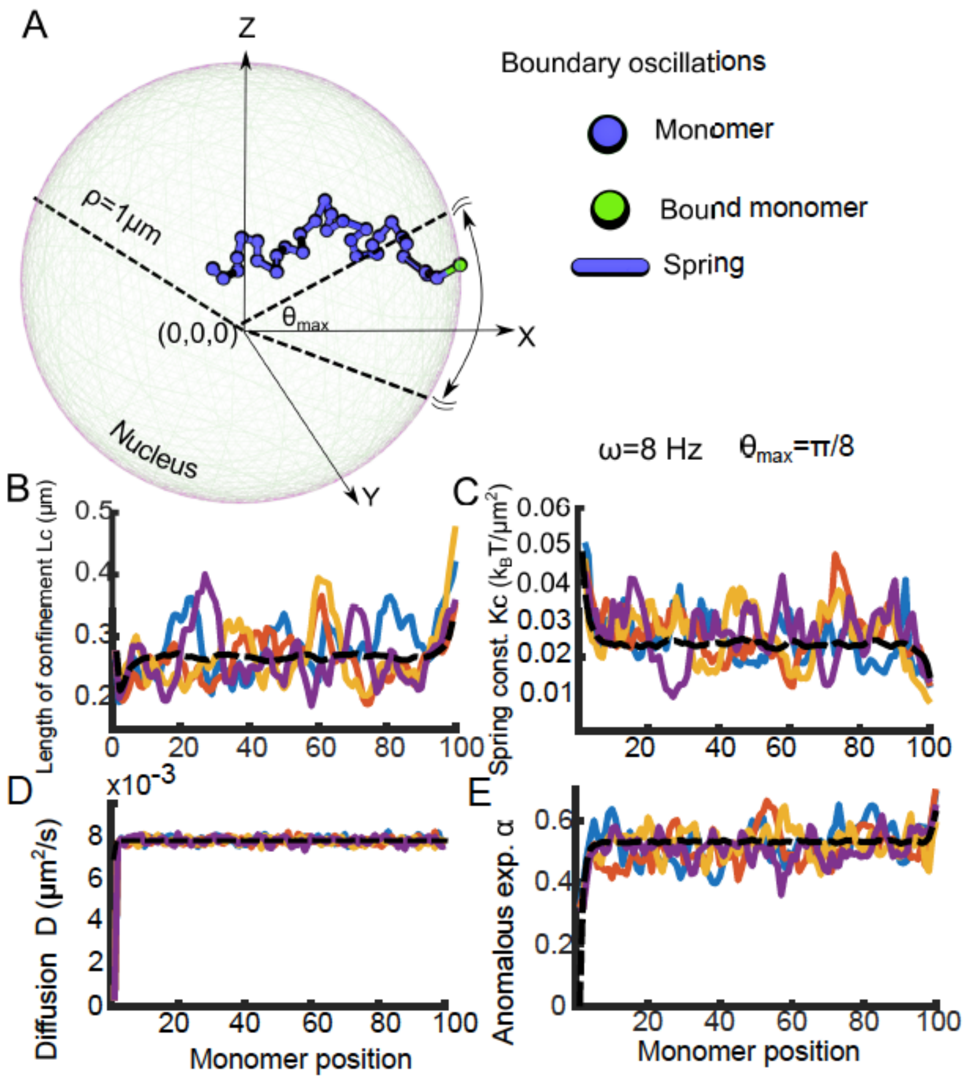

A
Z
Boundary oscillations
Monomer
Bound monomer
Spring
ρ=1μm
θmax
(0,0,0)
X
Y
Nucleus
ω=8 Hz θmax=π/8
B
Length of confinement Lc (μm)
C
Spring const. Kc (kBT/μm²)
D
x10-3
Diffusion D (μm²/s)
Monomer position
E
Anomalous exp. α
Monomer position

### Figure S4

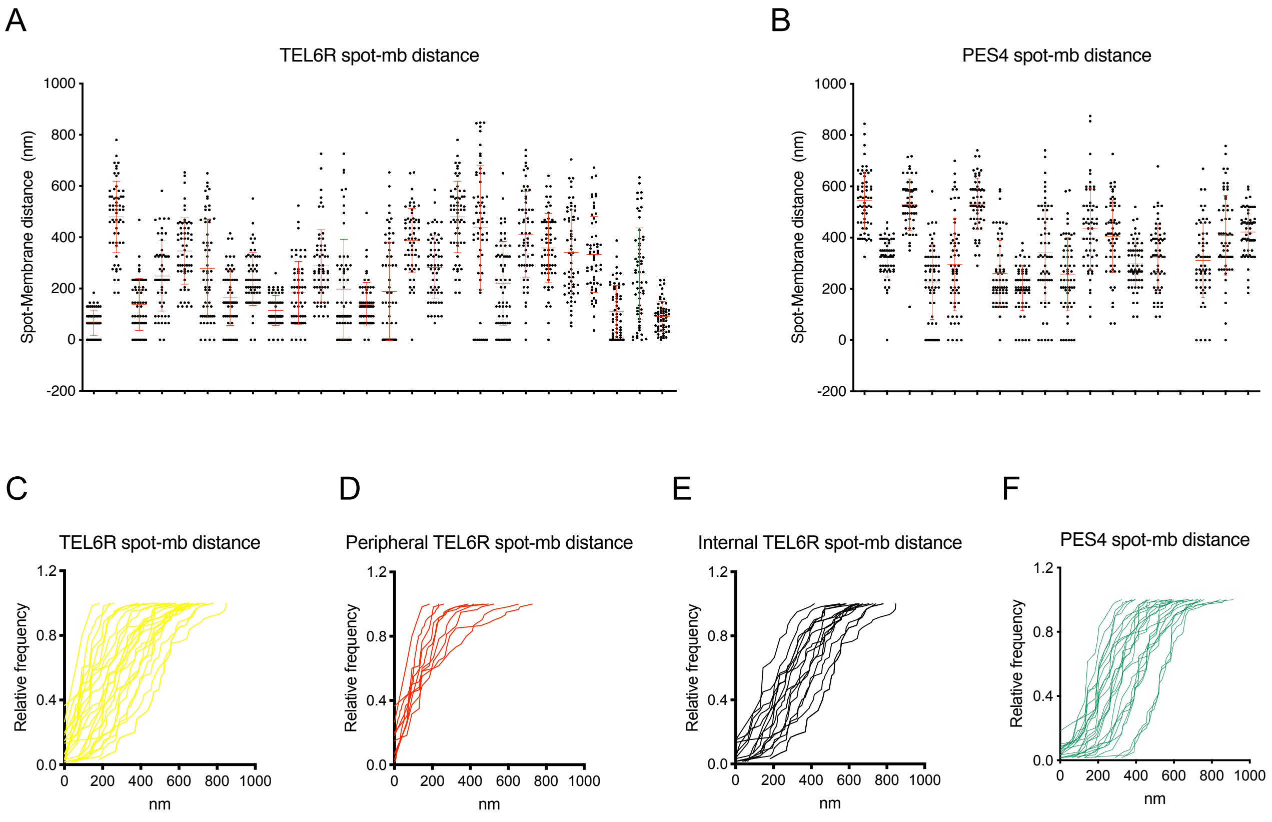
A
TEL6R spot-mb distance
Spot-Membrane distance (nm)
B
PES4 spot-mb distance
Spot-Membrane distance (nm)
C
TEL6R spot-mb distance
Relative frequency
nm
D
Peripheral TEL6R spot-mb distance
Relative frequency
nm
E
Internal TEL6R spot-mb distance
Relative frequency
nm
F
PES4 spot-mb distance
Relative frequency
nm

### Figure S5

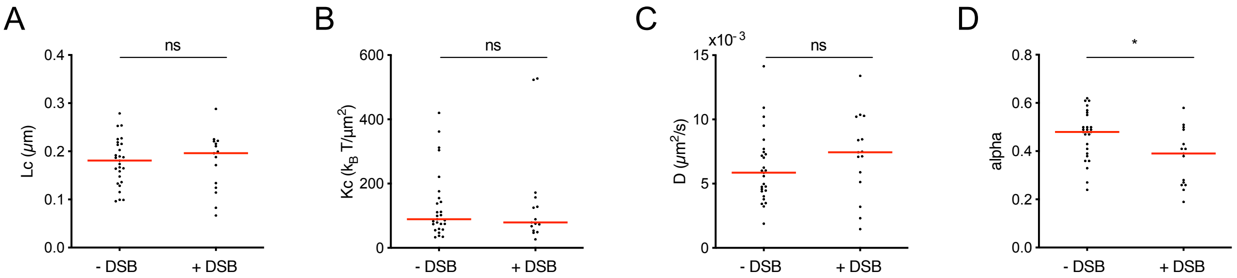

### Supplementary Methods

M. Toulouze A.Amitai O. Shukron, D. Holcman and K. Dubrana

April 11, 2019

### 1 Polymer models

#### 1.1 The RCL polymers

We now briefly describe the randomly cross-linked (RCL) polymer introduced in [2, 3] as a coarse-grained representation of the chromatin.

##### 1.1.1 Constructing the RCL polymer

We recall that a RCL polymer is composed of $N$ monomers located at position $\boldsymbol{R} = [r_1, r_2, ..., r_N]$, connected sequentially by spring constant, where $N_c$ added spring connectors between non nearest-neighboring randomly chosen monomers. A given configuration polymer with added monomer is a realization $\mathcal{G}$. The total energy of the RCL polymer is the sum of the spring potential of the linear backbone and that of the random connectors, given by

$$\phi(\boldsymbol{R}) = \frac{\kappa}{2}\boldsymbol{R}^T(\boldsymbol{M} + \boldsymbol{B}^{\mathcal{G}}(\xi))\boldsymbol{R}, \tag{1}$$

where $\boldsymbol{M}$ is the Rouse matrix for the linear backbone

$$\boldsymbol{M}_{m,n} = \begin{cases} -1, & |m-n| = 1; \\ -\sum\limits_{j\neq m} \boldsymbol{M}_{m,j}, & m = n; \\ 0, & \text{else}, \end{cases} \tag{2}$$

and $\boldsymbol{B}^{\mathcal{G}}(\xi)$ is the added connectivity matrix

$$\boldsymbol{B}^{\mathcal{G}}_{m,n}(\xi) = \begin{cases} -1, & |m-n| > 1 \; r_m \text{ and } r_n \text{ are connected;} \\ -\sum_{j \neq m} \boldsymbol{B}^{\mathcal{G}}_{m,j}(\xi), & m = n; \\ 0, & \text{othweewise.} \end{cases} \tag{3}$$

The connectivity fraction $\xi$ is related to the number of added connectors by the relation

$$\xi = \frac{2N_c}{(N-1)(N-2)}. \tag{4}$$

The choice of $N_c$ monomer pairs to connect is randomized over each realization $\mathcal{G}$ of the polymer. The added connectors can represent binding molecules such as CTCF and cohesin and their random position represents the local heterogeneous chromatin organization. The steady-state and transient properties of the RCL polymer are discussed in [2, 3].

The dynamics of monomers $\boldsymbol{R}$ is driven by the field of force derived from the energy 1 and that of random fluctuations, described by the Smoluchowski limit of the Langevin equation

$$\frac{\mathrm{d}\boldsymbol{R}}{\mathrm{d}t} = -\frac{dD}{b^2}\left(\boldsymbol{M} + \boldsymbol{B}^{\mathcal{G}}(\xi)\right)\boldsymbol{R} + \sqrt{2D}\frac{\mathrm{d}\boldsymbol{\omega}}{\mathrm{d}t}, \tag{5}$$

where $D$ is the diffusion coefficient, $b$ is the standard deviation of connectors' length, and $\boldsymbol{\omega}$ are $N \times d$ standard Brownian motions.

##### 1.1.2 Numerical simulations of the RCL polymer in spherical domain

Numerical simulations of Eq. 5 are performed in a spherical domain of radius $\rho = 1\mu m$, where each monomer is reflected. In each realization $\mathcal{G}$, we randomize the position of $Nc$ additional connectors. Each realization is simulated until the slowest polymer's relaxation time $\tau$ [2] that can be related to the time step $\Delta t$ and is defined by

$$\tau = \frac{b^2}{2D\Delta t(N\xi + 4(1-\xi))\sin(\pi/2N)}, \tag{6}$$

where $\xi$ is the connectivity fraction defined in 4, $N$ is the number of monomers. After $\tau$ steps, we continue simulations for an additional 3000 steps at time step $\Delta t = 0.033$, to reach a total simulation time of 100s. Simulation parameters are summarized in table 1.

| parameter | value [units] | description |
|---|---|---|
| $N$ | 32 or 64 or 100 | Number of monomers |
| $b$ | 0.2 ($\mu m$) | STD of connectors length |
| $D$ | $8 \times 10^{-3}$ ($\mu m/s^2$) | Diffusion coefficient |
| $T$ | 60 (s) | Simulation time |
| $\Delta t$ | 0.033 ($s$) | Simulation time step |
| $\Delta t_s$ | 0.33 ($s$) | Sampling time step |
| $\rho$ | 1 ($\mu m$) | Domain radius |
| $\theta_{max}$ | $[0, \frac{\pi}{256}, \frac{\pi}{128}, \frac{\pi}{64} \ldots, \pi]$ (rad) | Opening angle (rad) |
| $\omega$ | $\frac{2\pi}{T} \times [0, 0.5, 1, 2, 4, \ldots, 128]$ (Hz) | Oscillation frequency |
| $\sigma$ | 9 (monomers) | STD of connectivity distribution |

Table 1: parameters of the simulations with RCL polymer in a spherical domain.

#### 1.2 Simulation of a tethered Rouse polymer in an oscillating boundary

We simulated a Rouse chain with $N = 100$ monomers during $T = 60s$, where we fix the first monomer $r_1$ to the boundary of a spherical domain of radius $\rho = 1\mu m$ (see Fig. S2 and S3). To induce boundary oscillations, we restrict the tethered monomer position $r_1(t) = (x_1(t), y_1(t), z_1(t)))$ to oscillate on the $X - Z$ plane, with an amplitude $\rho \sin(\theta_{max})$ and frequency $2\pi\omega/T$, according to the equation

$$\begin{aligned} x_1(t) &= \frac{\rho \cos(\theta_{max})}{\sqrt{\cos^2(\theta_{max}) + \sin^2(\theta_{max}) \sin(2\pi\omega t/T)}}; \\ y_1(t) &= 0; \\ z_1(t) &= \frac{\rho \sin(2\pi\omega t/T) \sin(\theta_{max})}{\sqrt{\cos^2(\theta_{max}) + \sin^2(\theta_{max}) \sin(2\pi\omega t/T)}}. \end{aligned} \tag{7}$$

All other monomers $r_2 - r_{100}$ follows Rouse dynamics described in eq. 5, with $\boldsymbol{B} = 0$. We vary 1) the oscillation frequencies using ten values of $\omega \in \frac{2\pi}{T} \times [0, 0.5, 1.0, 2.0, 4.0, \ldots, 256]$, and 2) the oscillation amplitude using ten values of $\theta_{max} \in [0, \frac{\pi}{256}, \frac{\pi}{128}, \ldots, \pi]$.

### 2 Statistics of monomers of the RCL polymer in spherical domain

We describe now the four biophysical parameters that we are using to quantify the chromatin dynamics: these parameters are the anomalous exponent $\alpha$, effective diffusion $D_c$, length of constraint $L_c$, and the tethering constant $k_c$. These parameters are computed from trajectories of the RCL polymer generated for 100s post relaxation.

#### 2.1 Mean-squared-displacement of monomers $r_n$

The mean-square-displacement (MSD) of a monomer $r_n$ is defined as the average over realizations:

$$MSD_n(t) = \langle (r_n(t) - r_n(0))^2 \rangle. \tag{8}$$

For each monomer $r_n$ ($n = 1 \dots N$), we fitted the MSD (Eq. 8) using the function

$$f_{fitt}(t) = \beta_n t^{\alpha_n}, \tag{9}$$

where $\beta_n$ is a constant and the anomalous exponent of monomer $r_n$ is $\alpha_n$.

#### 2.2 Length of confinement $L_c$ of monomer $r_n$

The length of confinement $L_c(r_n)$ of monomer $r_n$ is computed by averaging over $N_s$ simulation steps, given by the empirical estimator [1]

$$L_c(r_n) = \sqrt{var(r_n)} \approx \sqrt{\frac{1}{N_s}\sum_{k=1}^{Ns}(r_n(k\Delta t) - \langle r_n \rangle)^2}, \tag{10}$$

where $\langle r_n \rangle$ is the mean position of monomer $n$.

#### 2.3 Effective spring coefficient $k_c$

We now recall how the spring parameter $k_c$ due to a resulting tethering force can be recovered from SPTs. A harmonic well of strength $k_w$ acting on a single monomer $\boldsymbol{r}_n$ of a polymer model [1] has an energy

$$U(\boldsymbol{r}_n) = \frac{1}{2}k_w(\boldsymbol{r}_n - \boldsymbol{\mu})^2, \tag{11}$$

where $\boldsymbol{\mu}$ is a fixed position for the center of the interacting force. The dynamics of an observed monomer $R_c$, can be recovered from many trajectories, because it is driven by this interacting force. The relation between the observed velocity extracted from SPTs and the tethering force is given by the following relation [1]

$$\lim_{\Delta t \to 0} \mathbb{E}\{\frac{\boldsymbol{r}_c(t+\Delta t) - \boldsymbol{r}_c(t)}{\Delta t}|\boldsymbol{r}_c(t) = \boldsymbol{x}\} = \ -Dk_{cn}\left(\boldsymbol{x} - \boldsymbol{\mu}\right), \tag{12}$$

where $\boldsymbol{r}_c(t)$ is the position of locus $c$ at time $t$ and $D$ the diffusion coefficient. $\mathbb{E}\{.|\boldsymbol{r}_c = \boldsymbol{x}\}$ is the conditional averaging, restricted to the condition that the tagged monomer is at position $\boldsymbol{r}_c = \boldsymbol{x}$. Relation (12) links the average velocity of the observed monomer $R_c$ to the force applied at a distance $|c-n|$. For a Rouse polymer with a potential well of type (11), the effective spring coefficient is given by

$$k_{cn} = \frac{k\kappa}{\kappa + |c-n|k}, \tag{13}$$

where $\kappa$ is the monomer-monomer spring coefficient. For a general polymer, relation 13 is not satisfied, and no general formula exists. The string constant $k_c$ can be estimated from relation 12 using the empirical estimator applied to the trajectories of the locus $\boldsymbol{r}_c(t)$ by

$$k_c \approx \frac{1}{2(N_p-1)} \sum_{i=1}^{2} \sum_{h=1}^{N_p-1} \frac{r_c^i((h+1)\Delta t) - r_c^i(h\Delta t)}{D_c \Delta t(r_c^i(h\Delta t) - \langle r_c^i \rangle)}, \tag{14}$$

where $i$ is the spatial dimension (in two dimensions, we sum over the $x$ and $y$ components) and $N_p$ is the number of points in the trajectory. In practice [1], the quantity $\langle r_c^i \rangle$ is computed by averaging over the trajectory. The diffusion coefficient $D_c$ can be computed by using formula 15. To avoid numerical instabilities, we use a linear regression to estimate the slope for $r_c^i((h+1)\Delta t) - r_c^i(h\Delta t)$ versus $(r_c^i(h\Delta t) - \langle r_c^i \rangle)$.

#### 2.4 Apparent diffusion coefficient $D_c$

The apparent diffusion coefficient $D_c$ of monomer $R_n$ is estimated by the empirical estimator

$$D(r_n) = \frac{1}{2dN_p\Delta t} \sum_{i=1}^{d} \sum_{k=1}^{N_p-1} (r_m((k+1)\Delta t) - r_n(k\Delta t))^2. \tag{15}$$

For monomers restricted to move only on the boundary of a domain, we set $d = 2$ in formula 15 to account for the motion on a two-dimensional manifold.

### 3 Simulation of a cross-link polymer model with a Gaussian connectivity distribution around the SPB

To account for the crowded motion of genomic loci located in the vicinity of the spindle pole body (SPB), we decided to add non-uniform cross-linkers so that genomic loci located far away from the SPB could be freely moving. To implement this scenario we distributed cross-linkers in a RCL model (see subsection 1.1) concentrated around the SPB. We model this connectivity distribution $f_{N/2}$ using a Normal distribution around the central monomers $r_{\frac{N}{2}}$ (the SPB), and given by

$$f_{\frac{N}{2}}(x) = \frac{1}{\sqrt{2\pi\sigma^2}} \exp\left(-\frac{x^2}{2\sigma^2}\right), \tag{16}$$

where $x$ is the genomic distance from the SPB and $\sigma$ is the standard-deviation. In each realization $\mathcal{G}$ of the polymer, we added $N_c$ connectors between monomer pairs $r_{i_k}, r_{j_k}$ randomly chosen, such that $\mathcal{G} = \{(i_1, j_1), (i_2, j_2), \ldots, (i_{N_c}, j_{N_c})\}$ are indices pairs sampled from $f_{N/2}$ by rounding sampled numbers to the nearest integer.

To account for the measured values of the anomalous exponent $\alpha$, the length of confinement $Lc$, the effective spring constant $Kc$ and the diffusion coefficient $D_c$, we restricted the RCL polymer with $N = 100$ monomers in a reflecting spherical domain of radius $\rho = 1\mu m$. We fixed monomer 50 to the boundary to represent the SPB, whereas monomers 1 and 100 can freely diffuse on the boundary of the domain. The simulation duration was 60s, with a time step $\Delta t = 0.003s$. To match the acquisition time intervals in experiments (Fig. 1), we sample trajectories at $\Delta t_s = 0.33s$ (i.e., every 100 simulation steps). Finally, we computed from simulation trajectories the values of $\alpha$ (Eq. 9), $D_c$ (Eq. 15), $Lc$ (Eq. 10),and $Kc$ (Eq. 13) for monomers 1-100. We then average the values of the four statistical parameters computed over simulations and compare them to the experimental ones. Finally, we adjusted the values of the polymer parameters: $b$, $D$, $\sigma$, and $Nc$, from simulations to obtain an optimal fit (in least square sense) to the experimental data of the four statistical parameters (Eq. 9-15). We obtain the following parameter values: number of connectors: $N_c = 60$, the standard deviation for the monomer distance is $b = 0.38\mu m$ and the standard deviation for the connectivity is $\sigma = 9$ (in monomer units). The rest of the parameters can be found in table 1. The results are shown in Fig. 5.
